# A Proteome-Wide Map of Chaperone-Assisted Protein Refolding in the Cytosol

**DOI:** 10.1101/2021.11.20.469408

**Authors:** Philip To, Yingzi Xia, Taylor Devlin, Karen G. Fleming, Stephen D. Fried

## Abstract

The journey by which proteins navigate their energy landscapes to their native structures is complex, involving (and sometimes requiring) many cellular factors and processes operating in partnership with a given polypeptide chain’s intrinsic energy landscape. The cytosolic environment and its complement of chaperones play critical roles in granting proteins safe passage to their native states; however, the complexity of this medium has generally precluded biophysical techniques from interrogating protein folding under cellular-like conditions for single proteins, let alone entire proteomes. Here, we develop a limited-proteolysis mass spectrometry approach paired with an isotope-labeling strategy to globally monitor the structures of refolding *E. coli* proteins in the cytosolic medium and with the chaperones, GroEL/ES (Hsp60) and DnaK/DnaJ/GrpE (Hsp70/40). GroEL can refold the majority (85%) of the *E. coli* proteins for which we have data, and is particularly important for restoring acidic proteins and proteins with three to five domains, trends that come to light because our assay measures the structural outcome of the refolding process itself, rather than indirect measures like binding or aggregation. For the most part, DnaK and GroEL refold a similar set of proteins, supporting the view that despite their vastly different structures, these two chaperones both unfold misfolded states, as one mechanism in common. Finally, we identify a cohort of proteins that are intransigent to being refolded with either chaperone. The data support a model in which chaperone-nonrefolders have evolved to fold efficiently once and only once, co-translationally, and remain kinetically trapped in their native conformations.

## INTRODUCTION

Protein folding represents the culmination of the central dogma of molecular biology – enabling the primary information encoded in nucleic acids and translated into polypeptides, to take shape into functional macromolecules. The striking accuracy of AI-based structure predictors has given new credence to Anfinsen’s dogma that protein three-dimensional structures is encoded at the amino acid sequence level (Anfinsen, 1973; Jumper et al., 2021); nevertheless, the journey by which proteins navigate their energy landscapes to locate their native structures is complex, involving (and sometimes requiring) many cellular processes and factors (Balchin et al., 2016; Tyedmers et al. 2010). Whilst it is well understood that molecular chaperones are required for specific proteins to refold from denatured forms (Brinker et al., 2001; Kerner et al., 2005; Singh et al., 2020; Viitanen et al., 1990), how these findings generalize to the proteome-scale is less clear; moreover, the potential influence of the cellular environment is typically not captured in most chaperone refolding experiments, leaving open questions about their physiological salience.

Traditional protein folding assays monitor structure or activity recovered by a denatured protein molecule following dilution from denaturant (Anfinsen, 1961); however, activity-based readouts are challenging to generalize to whole proteomes. Pioneering work by Kerner et al. introduced a high-throughput method to survey the clients of GroEL/GroES (*E. coli*’s group I chaperonin) by identifying proteins that are enriched in a fraction co-precipitating with chaperonin (Kerner et al., 2005), an approach that has since been extended to survey several other chaperone systems, such as DnaK (Calloni et al., 2012; Willmund et al., 2013). High-throughput measurements of protein precipitation, conducted on individually over-expressed proteins with and without chaperones (Niwa et al., 2009; Niwa et al., 2012), or on whole extracts following heat treatment (Mateus et al., 2018; Jarzab et al., 2020) have also been reported.

Despite these technical advances, systematically dissecting the roles that cellular processes and factors play in various proteins’ biogenesis remains challenging, even for the relatively simple *E. coli* proteome. Pull-down proteomic approaches cannot unambiguously assess a protein’s dependency (obligatory use) on a chaperone to refold, since the experiment measures association, rather than the structural outcome of the refolding process itself.

Indeed, many proteins that were presumed to be obligate chaperonin clients based on their enrichment on chaperonin co-precipitation studies were found to remain soluble *in vivo* during GroE knock-down (Fujiwara et al., 2010). Furthermore, a recent study (To et al., 2021) estimated that a third of soluble *E. coli* proteins are intrinsically nonrefoldable, meaning they cannot fully reassume their native forms following complete denaturation, even under conditions without appreciable precipitation. However, how many (and what kinds) of intrinsically nonrefoldable proteins can be rescued by chaperones – as opposed to requiring cotranslational folding (Fedorov and Baldwin 1997; Frydman et al., 1999; Liu et al., 2019) – is not known. A particularly underexplored question is when chaperones are required for refolding in the presence of the full complement of metabolites, ions, and small molecules that make up the cytosol.

To address these questions, we developed a limited-proteolysis mass spectrometry (LiP-MS) approach to probe protein structures globally during refolding (Figure 1, Figure 1– figure supplement 1A) (De Souza and Picotti, 2020; Feng et al., 2014; Park and Marqusee, 2005; Park et al., 2007). In this experiment, *E. coli* lysates are fully unfolded by overnight incubation in 6 M guanidinium chloride (GdmCl), returned to native conditions by rapid dilution, and the conformational ensembles of the proteins in the mixture probed by pulse proteolysis with proteinase K (PK), which cleaves only in regions that are solvent-exposed or flexible. Using liquid chromatography tandem mass spectrometry (LC-MS/MS), we sequence and quantify tens of thousands of peptide fragments to assess regions of proteolytic susceptibility and compare the proteolysis profile of each protein following refolding to that of its native form, recovered from the original lysate.

**Figure 1.**
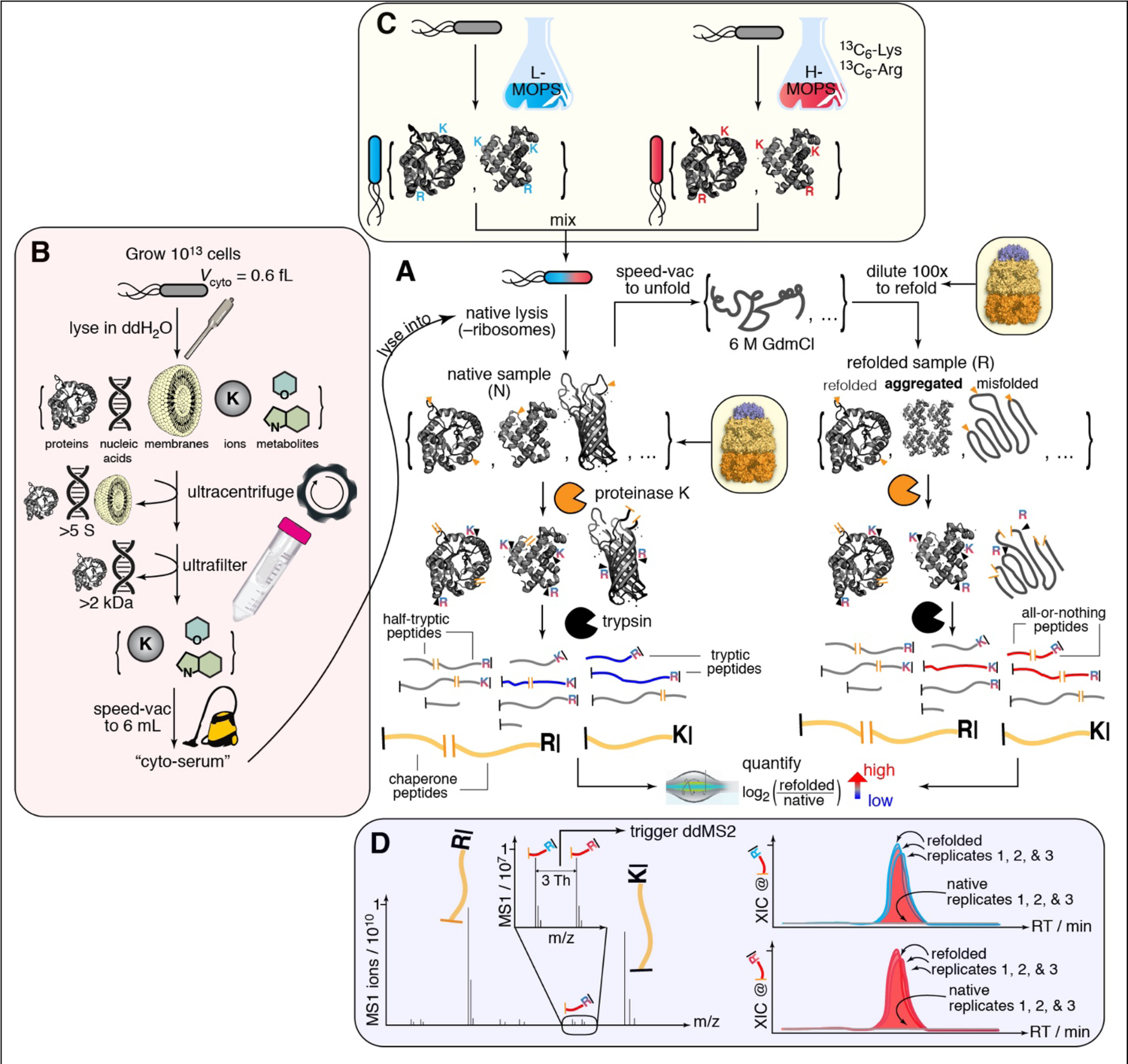
Limited-proteolysis mass spectrometry (LiP-MS) to interrogate the refoldability of the *E. coli* proteome under cellular-like conditions, highlighting methodological developments. (A) The core portion of the experiment. *E. coli* cells are lysed and ribosomes are depleted from the lysate. Extracts are globally unfolded and refolded by dilution. Proteinase K (PK) selectively cleaves at unstructured/exposed regions of proteins, thereby encoding structural information about the ensemble of protein conformations into cleavage sites. After quenching PK, proteins are fully trypsinized. The refolded/native abundance ratios are measured by label-free quantification (LFQ). (B) Preparation of cyto-serum. Large populations of *E. coli* cells are lysed by sonication into pure water, and all macromolecules are removed by ultracentrifugation and ultrafiltration. The liquid is reduced to the same volume that was enclosed by the original *E. coli* population’s cytoplasms. Cyto-serum is used as a lysis buffer for refolding experiments. (C) In chaperone refolding experiments, large concentrations of chaperone are supplemented into native samples and refolding reactions, which overwhelms the signal from proteome-derived peptides. To make proteome-derived peptides distinguishable, a pseudo-SILAC method is used in which replicate *E. coli* cultures are grown in rich MOPS media with either light (L) or heavy (H) lysine (Lys) and arginine (Arg). L/H pairs of cultures are mixed together and co-lysed. Consequentially, peptides derived from the proteome will exist as isotopomeric pairs. (D) Co-eluting isotopomer pairs are preferentially isolated for data-dependent MS2 (ddMS2) scans, enabling high coverage of the *E. coli* proteome during MS analysis. In label-free quantification (LFQ), peptide abundance ratios (refolded/native) are calculated by comparing the area under the curve of the extracted ion chromatograms (XIC) at the appropriate retention time (RT) across separately-injected samples. Non-refolding regions of proteins generate half-tryptic fragments that are absent in the native sample. Pseudo-SILAC provides greater confidence that the signal is absent in natives by doubling the number of quantifiable features for each peptide group.

Using this approach, we interrogate protein refolding under conditions that more closely recapitulate the cellular context, in the cytosol, with the molecular chaperones, GroEL/ES (Hsp60) and DnaK/DnaJ/GrpE (Hsp70/40). We discover that protein isoelectric point (pI) emerges unexpectedly as a key explanatory variable for refoldability: basic proteins are generally efficient refolders, particularly in the cytosolic milieu, whilst acidic proteins are more frequently reliant on GroEL to refold. GroEL can restore many intrinsically nonrefoldable proteins, especially acidic proteins, proteins with high molecular weight (MW), proteins with three to five domains, and domains with α/β architectures. The cohort of proteins that GroEL refolds overlaps extensively with those which DnaK can restore, suggesting a common mechanism for these two distinct molecular machines. Finally, our study sheds light on a small group of proteins that are recalcitrant to refolding with either chaperone, a group that most likely is adapted to fold co-translationally and remain trapped in their native states, obviating the need for chaperone assistance after their synthesis. This group heavily represents proteins involved in core and ancient metabolic processes, namely glycolysis and translation. Structural, mechanistic, and evolutionary implications of these ‘chaperone-nonrefolders’ are discussed.

## RESULTS

### A Method to Interrogate Refolding the *E. coli* Proteome in Cytosol with Chaperonin

The *E. coli* cytosol is an idiosyncratic medium predominantly buffered by glutamate and replete with a wide array of cofactors, metabolites, and ions with concentrations spanning over 6 orders of magnitude (Bennet et al., 2009). To probe the effect this medium exerts on protein folding, we isolate the cytosolic medium by culturing cells to the end of log phase and lysing them into pure water (Figure 1B). Macromolecules larger than 2 kDa are depleted by ultra-centrifugation and subsequent ultra-filtration of the supernatant (see Methods, Figure 1 –figure supplement 2A). The filtrate is then reduced under vacuum until its volume equals that of the combined internal volume of the original cellular population, given the estimated *E. coli* cytoplasm volume of 0.6 fL/cell (Philips et al., 2008). The resulting liquid, which we refer to as ‘cyto-serum,’ consists of all the stable and free ions, metabolites, and cofactors present in the *E. coli* cytosol near their physiological concentrations. Cyto-serum is a non-viscous off-yellow (λ_max_ 258 nm) liquid with a pH of 7.4 and an ATP concentration of 0.6 mM (Figure 1–figure supplement 2A-F).

We use cyto-serum as a lysis buffer to resuspend separate *E. coli* cell pellets (grown to the end of log phase in MOPS media (Neidhardt et al., 1974)), which are natively lysed by cryogenic pulverization, a mechanical lysis method chosen because it keeps large and weakly-bound protein assemblies intact (Harris, 1987; Wallace et al., 2015) (Figure 1). Use of cyto-serum as a lysis buffer enables us to maintain proteins at suitably low concentrations for refolding (0.116 mg/ml, ca. 4 µM), whilst keeping the small molecule constituents of the cytosol near their physiological concentrations.

In preliminary experiments, we tested whether cyto-serum would be suitable for global refolding experiments by measuring the levels of aggregation that accrue after 2 h. Pelleting assays detected low but non-zero levels of aggregation (6 ± 2% of protein), in contrast with the buffer conditions we had previously devised (20 mM Tris pH 8.2, 100 mM NaCl, 2 mM MgCl_2_, 1 mM DTT) that result in very low levels of precipitation (3 ± 1%, Figure 1–figure supplement 2D). This 3% increase in aggregation is close to what we previously observed for refolding in a defined buffer at neutral pH (To et al., 2021; Wang et al., 2017), thereby confirming that alkaline pH helps suppress aggregation, and that the cytosolic components do not increase aggregation levels beyond an expected effect from pH. To further investigate aggregation (including smaller soluble non-precipitating aggregation), we performed sedimentation velocity analytical ultracentrifugation (AUC) and mass photometry (MP) on these refolding reactions (Figure 1–figure supplement 3). Both techniques showed that the molecular size distributions of the refolded samples were similar to native extracts, confirming the absence of soluble aggregates. Overall, these studies suggest that complex mixtures of proteins are less aggregation-prone than most of these individual proteins are when they are overexpressed (e.g., Niwa et al., 2009).

Following these tests, we proceeded to perform global refolding experiments by diluting unfolded *E. coli* extracts with cyto-serum supplemented with 4 µM GroEL and 8 µM GroES (Figure 1; ca. 100-fold higher concentration than their natural abundances in diluted lysate). Because it is important to compare compositionally identical native and refolded samples, we also supplement chaperones and cyto-serum into the native samples and equilibrate them together for 90 min prior to limited proteolysis (cf. Figure 1A). This step is essential because even though native proteins shouldn’t “need” GroEL, if a correctly-refolded protein has a propensity to associate transiently with GroEL (as a “triage complex” (Gottesman et al., 1997; Powers et al., 2012)), such an interaction would still affect its proteolysis profile and therefore needs to be present in the native sample.

In preliminary LC-MS/MS experiments, we detected low coverage of the proteome because >80% of the total protein content in these refolding reactions are the added chaperone and cyto-serum adds many non-protein contaminants (Figure 2A). To address this challenge, we developed an isotope-labeling strategy to distinguish peptides belonging to refolding clients from those belonging to chaperonin proteins or from other cellular contaminants (Figure 1C). Three replicate *E. coli* cultures are grown in two different MOPS media: one with natural abundance (light) isotopes of Arg and Lys, and a second with [^13^C_6_]Arg and [^13^C_6_]Lys (heavy). Pairs of light and heavy media are mixed together (for each biological replicate) prior to lysis and initiating the unfolding/refolding/LiP-MS workflow. In this way, peptides from client proteins will be present in the sample as a pair of isotopomers that co-elute during liquid chromatography and generate a signature twin-peak feature (Figure 1D) that distinguish them from chaperone-derived peptides despite being several orders of magnitude lower in intensity (Figure 2C). The mass spectrometer is then instructed to preferentially select peaks with the correct spacing for data-dependent isolation and MS2 acquisition. We confirmed that co-eluting isotopomers generate fragmentation spectra with expected mass-shifts in the y-ions (Figure 2D).

**Figure 2.**
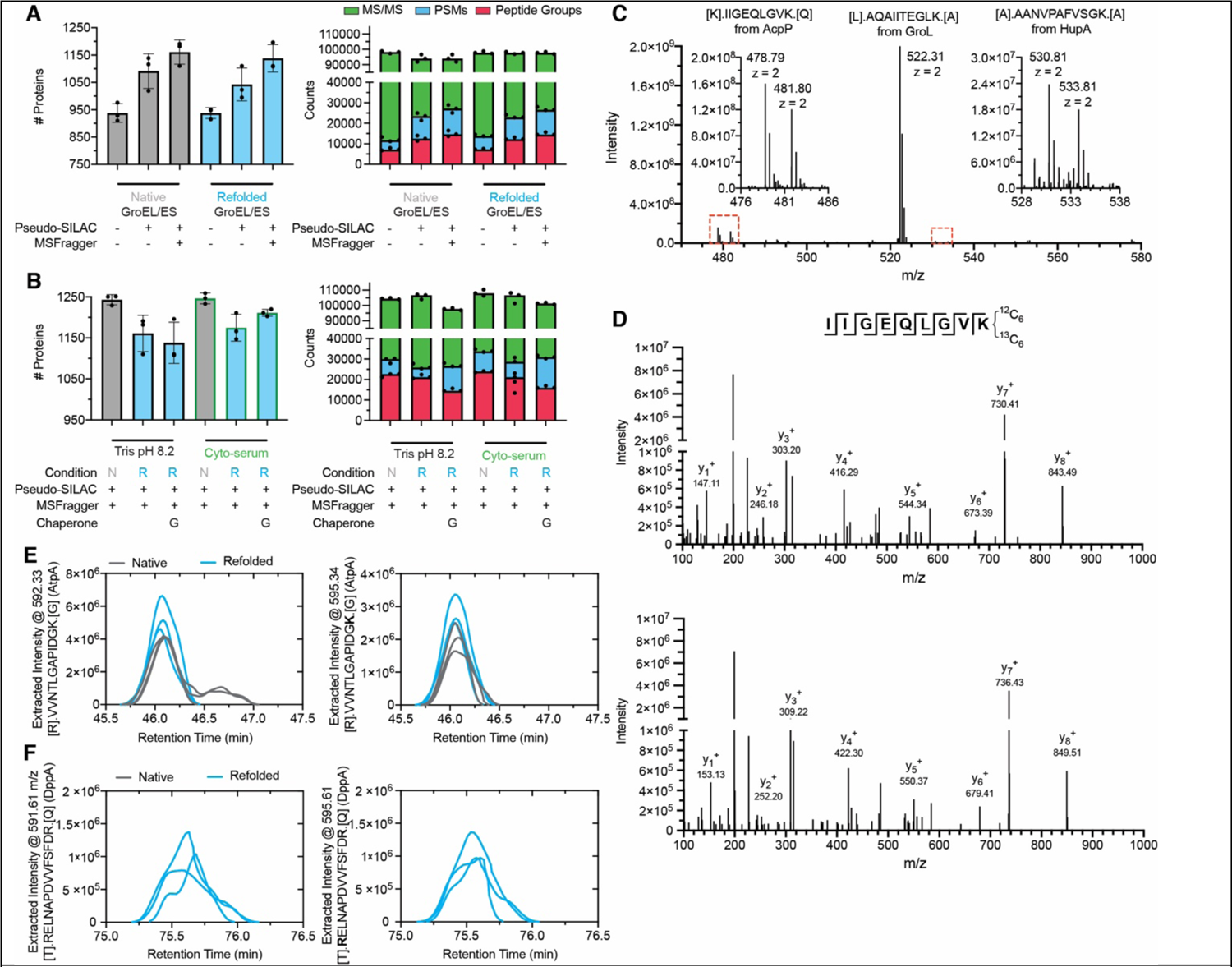
Pseudo-SILAC to distinguish extract-derived peptides from chaperone-derived peptides. (A) Bar charts showing the number of proteins and peptide groups identified, peptide-spectrum matches (PSM), and total MS/MS spectra obtained on individual runs, testing the effect of the pseudo-SILAC method and the MSFragger search algorithm. The number of MS/MS spectra acquired per run is similar, but pseudo-SILAC increases the number of unique PSMs and peptide groups by selecting features that are *not* chaperonin or cytosolic contaminants for data-dependent MS2 acquisition. (B) Bar charts showing that the combination of pseudo-SILAC and MSFragger results in no significant loss in coverage in experiments conducted in cyto-serum and with GroEL/ES. (C) A sample MS1 spectrum from a refolding experiment with GroEL/ES in cyto-serum. Peptides derived from refolded proteins, but not from chaperone, display twin-peaks separated by 3 Th (as expected for a doubly-charged peptide). (D) Sample MS2 fragmentation spectra from two co-eluting peptides that differ only by the isotopic composition of the C-terminal lysine. The y-ions (indicated) are all displaced by 6 Th, as expected for singly-charged fragments. (E) Extracted ion chromatograms for the peptide indicated (from AtpA) in three replicate native samples and in three replicate refolded samples, at two m/z’s corresponding to the light and heavy-substituted isotopes. The abundance of the peptide is similar in the native and refolded forms, in both isotope states, implying that this region of AtpA properly refolded; i.e., had the same PK susceptibility in both forms. (F) Similar to panel E but for an all-or-nothing peptide from DppA which is not detected in any of the three native replicates at the m/z for both the light- and heavy-substituted isotope.

Combining this method with the MSFragger spectral search algorithm (Kong et al., 2017), we were able to reproducibly obtain satisfactory numbers of unique identifications and quantifications (Figure 2A-B). Moreover, this approach enables us to double the number of independent quantifications used to assess the refolded/native ratio for many peptides, providing additional confidence in our measurements (Figure 2E-F). This verification is particularly salient for verifying so-called “all-or-nothing peptides” which are detected exclusively in the refolded (or native) samples (Figure 2F) and provide compelling evidence for a structural difference in the refolded form of a protein. Not detecting a feature across replicates in two distinct mass channels provides further weight to the assertion that missing spectral features constitute *evidence of absence* (Figure 1D). We refer to this strategy as ‘pseudo-SILAC’ because it uses stable isotope labeling to direct the mass spectrometer to select the correct features, as opposed to performing quantifications. Instead, we calculate refolded/native abundance ratios by comparing the areas under the curve between runs (known as label free quantification (LFQ)), because of its superior dynamic range (Palomba et al., 2021; Nahnsen et al., 2013) and ability to confidently identify when a feature is *absent* from a particular sample. We note that even though pseudo-SILAC is not as necessary for experiments without chaperones, we applied it to all conditions in this study uniformly to remove any potential source of bias when comparing chaperone to non-chaperone conditions.

### GroEL/GroES Rescues Many Nonrefoldable Proteins

GroEL/GroES significantly remodels the refolding profile of the *E. coli* proteome (Figure 3). To summarize these data, we present peptide-level volcano plots and abundance ratio histograms (Figure 3A-B) for refolding in cyto-serum without and with chaperonin after 1 min, where the differences are the most apparent. Half-tryptic peptides are shown in blue, and demarcate locations where PK cleaved (cf. Figure 1); full-tryptic peptides are shown in black, and represent the absence of a PK cut. The observation that most peptides that are more abundant in the refolded samples (right-hand side) are half-tryptic (86% without GroEL, 90% with GroEL), and that most peptides that are more abundant in the native samples (left-hand side) are full-tryptic (80% without GroEL, 81% with GroEL; P < 10^-15^ by Mann-Whitney U-test for both) imply that the refolded proteome is globally more susceptible to proteolysis than the native proteome.

**Figure 3.**
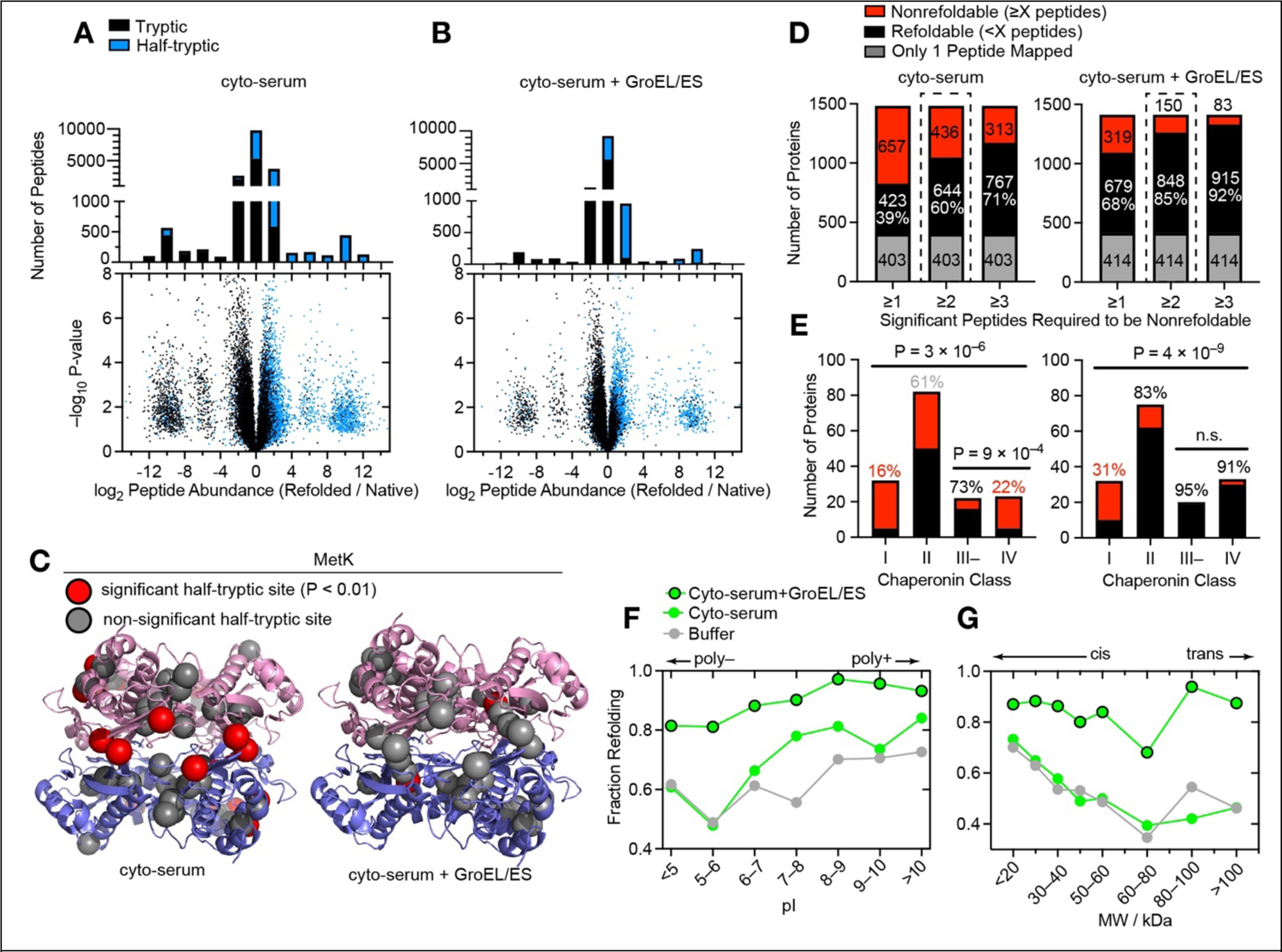
GroEL/ES is a versatile chaperone that assists the refolding of many *E. coli* proteins. (A) Volcano plots and associated peptide histogram comparing peptide abundances from 3 native and 3 refolded *E. coli* lysates after 1 min of refolding for experiments in which cells were lysed in cyto-serum (natives), or lysed in cyto-serum, unfolded overnight in 6 M GdmCl, and refolded in cyto-serum for 1 min. Effect sizes reported as ratio of averages, and P-values are calculated using the *t* test with Welch’s correction for unequal variance (*n* = 3). “All or nothing” peptides form the two lobes centered at ±10 of the abscissa, and in order to be considered they had to be detected in all 3 replicates of one sample-type (refolded or native) and zero out of 3 of the other. Half-tryptic peptides are denoted in blue, tryptic peptides in black. Data correspond to #1 in Figure 1–figure supplement 1B. (B) Similar to panel A, except where 4 µM GroEL and 8 µM GroES were present in the native samples and added to the refolding reaction. Data correspond to #4 in Figure 1–figure supplement 1B. (C) Structure of MetK (PDB: 1P7L), indicating sites where proteolytic susceptibility is the same (gray spheres) or significantly different (red spheres) in the refolded samples compared to native. Left, locations of 9 PK cut-sites with significantly different susceptibility in the refolded sample, after refolding in cyto-serum (red spheres). Right, location of one PK cut-site with significantly different susceptibility in the refolded sample, after refolding cyto-serum and GroEL/ES. (D) Bar charts showing the total number of proteins assessed, of which how many are designated refoldable or nonrefoldable for refolding experiments in cyto-serum, without and with GroEL/ES. Bars correspond to alternative cutoff schemes, varying the number of peptides with significant difference in proteolytic susceptibility in refolded form required to call a protein nonrefoldable. ≥2 is used for the rest of the study. In gray are proteins with only 1 peptide quantified, which are not used in further analyses (and not counted in the refolding percentages). Data correspond to #2 and #5 in Figure 1–figure supplement 1B. (E) Bar charts indicating the number of refolding and nonrefolding proteins associated with one of four chaperonin classes (as defined by Kerner et al., 2005; Fujiwara et al., 2010), in experiments without and with chaperonin. Percents indicate percentage refolding within that category. P-values for the all-way comparison are from chi-square test; for the two-way III– v. IV comparison are from Fisher’s exact test. (F) Fraction of proteins that refold in either Tris buffer (gray (Nissley et al., 2021)), cyto-serum (green), or cyto-serum with GroEL/ES (green, black border), separated on the basis of individual proteins’ isoelectric point (pI). Data correspond to #2 and #5 in Figure 1–figure supplement 1B. (G) Fraction of proteins that refold in either Tris buffer (gray (Nissley et al., 2021)), cyto-serum (green), or cyto-serum with GroEL/ES (green, black border), separated on the basis of individual proteins’ molecular weight (MW).

Points on the flanking lobes correspond to peptides that were detected only in the refolded or native samples. We refer to these as ‘all-or-nothing’ peptides and assign a limit-of-detection abundance to them in samples where they’re not detected. All-or-nothing peptides represent nonrefoldable regions within proteins that were *completely* inaccessible to PK in the native conformation but became proteolytically susceptible when that region failed to refold. After refolding with GroEL, many fewer all-or-nothing peptides were detected (1736 (9.5%) without GroEL, 691 (5.6%) with GroEL), signifying fewer proteins that were structurally distinct from their native forms. The number of all-or-nothing peptides decreases following 5 min of refolding, though the trend with respect to chaperonin is the same (1276 (7.6%) without GroEL, 515 (4.1%) with GroEL). Here we note that we employ a stringent filtering process to assess peptides with missing data (see Methods) and are confident in their status based on high reproducibility over nine orders of magnitude across technical replicates of the experiment (Figure 3–figure supplement 1).

We mapped peptides back to their parent proteins and labeled an individual protein nonrefoldable if we could identify two or more peptides with a significant abundance difference in the refolded samples relative to the native samples (>2-fold effect-size, P < 0.01 by t-test with Welch’s correction for unequal population variances). For MetK, there is only one such significant peptide after refolding with GroEL (Figure 3C) – many fewer than after refolding on its own – consistent with its known status as an obligate GroEL client (Ying et al., 2005). By this metric, the proteome was the most refoldable at the 5 min timepoint both with and without chaperonin (Figure 3–figure supplement 2A), hence we chose to focus on it for further analysis. After 5 min, in cyto-serum 60% of 1080 proteins are refoldable intrinsically (Data S1), and with the addition of GroEL/GroES, this rises to 85% of 998 proteins (Figure 3D, Data S2), using a ≥2 peptide cutoff to call a protein nonrefoldable (as used previously (To et al., 2021)). The overall refoldability rates do depend on this admittedly arbitrary cutoff employed to call a protein nonrefoldable; however, the ≥2 peptide cutoff can be viewed as a compromise between not allowing too much weight to be assigned to a single significant peptide, and not making it too difficult to call a protein nonrefoldable with lower coverage. Importantly, none of the key trends we describe in the following depend sensitively on this choice (Figure 3D, Figure 3-figure supplement 3G-I).

To contextualize this experiment, we first sought to compare these results to two landmark studies interrogating *E. coli* chaperonin usage across the proteome. Kerner et al. (2005) formalized a classification system based on the enrichment level of various proteins in the fraction that co-precipitates with a tagged GroEL/ES complex. Class I proteins are those that are de-enriched in the GroEL fraction relative to their level in the cytoplasm, whilst class III proteins are those that are highly enriched in the GroEL fraction. Complementing this study, Fujiwara et al. (2010) used an *E. coli* strain in which GroEL expression is arabinose dependent and measured which proteins precipitate in the *E. coli* cytoplasm after GroEL expression is cut off by shifting cells from arabinose to glucose. Many (40%) of the class III proteins were still soluble in the cytoplasm without chaperonin and were renamed class III^-^. On the other hand, those whose solubility in the cytoplasm is expressly chaperonin-dependent were renamed class IV.

Our refolding assay is strikingly consistent with Fujiwara’s sub-classification (Figure 3E). In the chaperonin-null condition (Figure 3E), the majority (73%) of class III^-^ proteins are refoldable, whereas only a minority (22%) of class IV proteins are. The observation concerning class III^-^ proteins implies, intriguingly, that there are many proteins that associate strongly with GroEL *in vivo* that do not actually require it. The strong alignment between class IV and nonrefoldability implies that most class IV proteins populate misfolded states which aggregate at the high concentrations of the cellular environment, but in our assay instead persist as soluble misfolded states that do not aggregate but also *cannot* correct themselves. The observation that a few class IV proteins are refoldable in our assay suggests that in these situations, GroEL’s function is to serve as an obligatory holdase, a function that is no longer necessary when aggregation is suppressed. With chaperonin added to the refolding reactions, both class III^-^ and class IV proteins are nearly completely refoldable (95% and 91% respectively, Figure 3E). This finding implies that the majority of GroEL’s obligate clients (class IV) require it *actively* (e.g., either as a foldase or unfoldase), not merely as an infinite-dilution chamber (e.g., holdase) (Brinker et al., 2001). It also shows that most obligate GroEL clients do not require additional chaperones in a hand-off mechanism (Langer et al., 1992). We note that class IV proteins are actually more refoldable in Tris buffer pH 8.2 (which further suppresses aggregation) than in cyto-serum (Nissley et al., 2021), hence in the cytosol, GroEL’s assistance is even *more* needed (because of greater aggregation propensity) than it is in an alkaline refolding buffer.

Proteins with higher isoelectric points (pI > 8) tend to be intrinsically refoldable and especially so in the cytosol, whereas proteins with lower isoelectric points (pI < 7) are less intrinsically refoldable, a difference that is largely mitigated by GroEL (Figure 3F). Proteins with high molecular weight (MW) tend to be less intrinsically refoldable, but GroEL smooths over this difference as well (with an important exception for proteins sized 60–80 kDa), exerting its most prominent rescuing power on proteins of greatest molecular weight (Figure 3G). The discontinuity for proteins sized 60–80 kDa has previously been attributed to the dimensions of the GroEL cavity, which is known not to accommodate proteins larger than 60 kDa (Kener et al., 2005). However, we find that GroEL is extremely effective at assisting the largest *E. coli* proteins. These observations support the theory that the unsealed *trans* cavity of GroEL is also an active chaperone, and are consistent with previous works that have found activity of GroEL on large substrates (Weissman et al., 1996; Chaudhuri et al. 2001; Chaudhuri et al. 2009; Farr et al., 2003; Paul et al., 2007).

Our data further elucidate the types of proteins that tend to be obligate GroEL refolders (Figure 4). To make this assessment, we pooled together the raw data from the experiments both with and without GroEL/ES, selected the subset of proteins that were confidently assessed in both conditions, and assigned them statuses based on their refolding outcomes in the two conditions (Figure 4A-B, Data SA). Inspection of the distribution of obligate GroEL refolders, broken down by pI range (Figure 4C), shows that obligate GroEL refoldability peaks for mildly acidic proteins (5 < pI < 6; 26%), is lower for proteins that are neutrally-charged in the cytosol (7 < pI < 8; 11%), and is lowest for basic proteins (pI > 10; 2%). Indeed, amongst polybasic proteins (pI > 10) there are three examples (7.3%) of proteins that lose their intrinsic capacity to refold in the presence of chaperonin (note only 1% of all proteins overall are in this category). This may be because some basic proteins could get stuck in the GroEL cavity, whose lumen is negatively charged (Tang et al., 2006; Tang et al., 2008). Such a tendency might explain why polybasic proteins generally have been optimized to refold on their own in the cytosol (Figures 3F, 4C), as they might otherwise unproductively bind too tightly within GroEL. As expected, low-MW proteins are the least likely to require GroEL, and high-MW proteins are the most (Figure 4D). The large (>80 kDa) obligate GroEL refolders are all (100%) multi-domain proteins, wherein potentially one non-native domain could fit in the unsealed *trans* cavity. Indeed, we find a robust trend that proteins with more domains up to 4 become progressively more reliant on GroEL (Figure 4E), though proteins with >5 domains appear to be poor refolders even with GroEL (P = 0.02 by chi-square test). Together, these findings provide support for the view that the *trans* mechanism is effective at resolving misfolded domains in the context of large multi-domain proteins.

**Figure 4.**
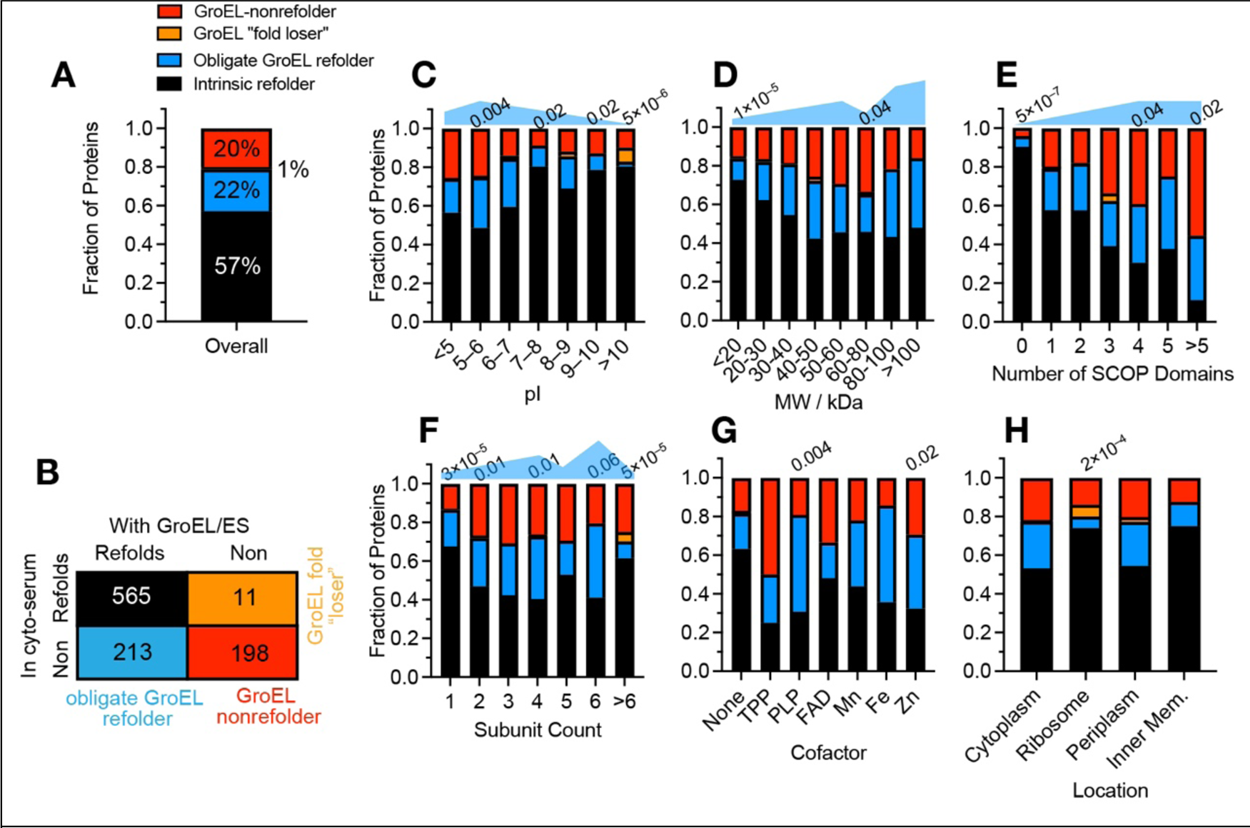
Defining the scope of obligate GroEL refolders across the proteome. (A) Frequency of proteins that refolded in both conditions (intrinsic refolder; black), only with GroEL/ES (obligate GroEL refolder; blue), only without GroEL/ES (GroEL “fold loser”; orange), or did not refold in either (GroEL-nonrefolder; red). Data correspond to #b in Figure 1–figure supplement 1C (Data SA). Numbers listed above bars indicate P-values by the chi-square test that the category has a different GroEL usage profile than the proteome overall. Blue shapes qualitatively denote the need-level for GroEL. (B) Truth table showing the number of proteins in each of the categories described in A. Analysis covers 987 proteins for which at least 2 peptides could be confidently quantified in both conditions. (C) As A, except proteins are separated on the basis of isoelectric point (pI). (D) As A, except proteins are separated on the basis of molecular weight (MW). (E) As A, except proteins are separated on the basis of the number of domains in the protein, as defined by the SCOP database. (F) As A, except proteins are separated on the basis of the number of subunits in the complex to which they are part. (G) As A, except proteins are separated on the basis of their bound cofactor. (H) As A, except proteins are separated by their cellular location.

We also found a few correlations between GroEL usage patterns and subunit composition, cellular location, and cofactors (Figures 4F–H). Monomers and assemblies of all sizes benefit from GroEL’s assistance, though with a noticeable dip at dimers and trimers, which is intriguing because these assemblies are the most likely to assemble co-translationally (Shiber et al., 2018; Shieh et al., 2015; Bertolini et al, 2021) (Figure 4F). Tetramers and hexamers are most likely to be obligate GroEL refolders (32% and 39% respectively), consistent with several model GroEL clients being tetramers like MetF (Singh et al., 2020) and DapA (Georgescauld et al., 2014; Ambrose et al., 2015). Proteins in large complexes with >6 subunits are the least reliant on GroEL (Figures 4F). We find that GroEL benefits cofactor-harbouring proteins, particularly proteins that host TPP, PLP, Fe^2+^, and Zn^2+^, which are generally less refoldable on their own (To et al., 2021), and have high propensities to be obligate GroEL refolders (between 38–50%, Figure 4G). Fe^2+^ and Zn^2+^ form strong near-covalent linkages with the coordinating residues Cys and His, hence, incorrect coordination would create an energetically entrenched misfolded state that might require energy input to undo. Finally, we find that GroEL is effective at recovering proteins in all *E. coli* locations (Figures 4H), including the periplasm. The observation is unusual because GroEL is strictly a cytosolic chaperone, and when extracted from cells does not co-precipitate periplasmic proteins (Kernet et al., 2005). Hence, even though periplasmic proteins use a distinct suite of chaperones *in vivo* (Mas et al., 2019), it would appear that GroEL can act as an effective substitute during *in vitro* refolding.

All of the trends that we have discussed here are statistically significant at the peptide level as well (P-values range from 10^-5^ to 10^-31^ by the chi-square test), implying that these differences in refoldability cannot be attributed to coverage bias (Figure 3–figure supplement 3A–F). Furthermore, the protein-level trends are not sensitive to the peptide cutoff to call a protein nonrefoldable (Figure 3–figure supplement 3G-I), and hence can be considered robust.

### Effect of Chaperonin on Refolding Kinetics

Classic protein folding kinetics studies, typically carried out on small single-domain proteins, record folding times on the ms–s timescales (Bartlett and Radford, 2009). Because of the duration of the PK incubation time (1 min), our experiments do not afford the same level of temporal resolution; however, comparisons between refoldability levels at the 1 min and 5 min timepoints can provide insight into the types of proteins that refold slowly (i.e., require more than 1 min) – both with and without chaperonin (Figure 5). In cyto-serum, overall refoldability increases from 52% to 60% from 1 to 5 min, a similar uptick as to what we observe in the chaperonin refolding experiment (77% to 85%). However, from 5 min to 2 h the overall refoldability in cyto-serum slightly decreases, which we attribute to a mix of degradation and aggregation (Figure 3–figure supplement 2A, B, E). With chaperonin, refolding decreases precipitously at 2 h (down to 74%), presumably due to depletion of ATP.

**Figure 5.**
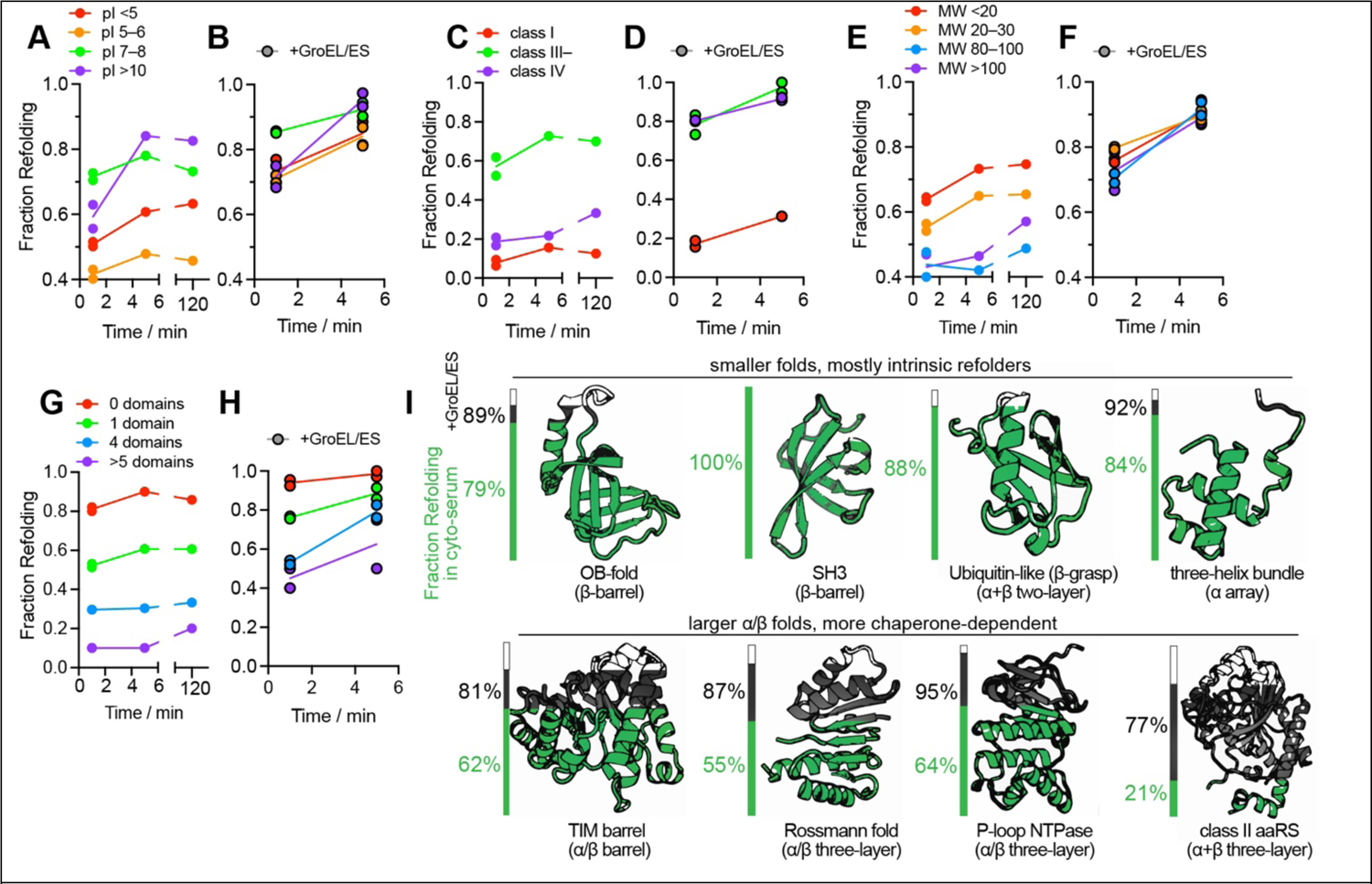
Kinetic of protein refolding in cyto-serum, with or without GroEL/ES. (A,B) Fraction of proteins that refold after 1 min, 5 min, or 120 min in either cyto-serum (A), or in cyto-serum with GroEL/ES (B, black borders), separated on the basis of individual proteins’ pI. Data correspond to #1-5 in Figure 1–figure supplement 1B. “Slow refolding” means requiring more than 1 min, but less than 5 min, to refold. “Very slow refolding” means requiring more than 5 min, but less than 2 h, to refold. Proteins with high pI are more likely to refold slowly both with and without GroEL. (C,D) As A and B, except proteins are separated on the basis of chaperonin class (Kerner et al., 2005; Fujiwara et al., 2010). (E,F) As A and B, except proteins are separated on the basis of molecular weight (MW). (G,H) As A and B, except proteins are separated on the basis of number of domains, as defined by the SCOP database. (I) Fraction of domains that refold in either cyto-serum (green) or cyto-serum with GroEL/ES (black), separated on the basis of which Fold the domain is assigned to in the SCOP hierarchy. Data correspond to #2 and #5 in Figure 1–figure supplement 1B. For these analyses, half-tryptic and tryptic peptides were mapped to the individual domains within a protein, based on residue ranges from alignments to HMMs.

Despite the similar increase in refoldability percentages from 1 to 5 min, the types of proteins that benefit from additional time were distinct without and with chaperonin. In the GroEL-null condition (Figure 5A, C, E, G, Data S1K), slow refolders tend to have high pI (>10; Figure 5A) or be class III^-^ (Figure 5C). These features are readily explainable: highly polycationic proteins would have significantly more intra-chain repulsion that would slow down compaction, and class III^-^ proteins are those which populate kinetically-trapped intermediates that, given time, can self-correct. Such proteins employ GroEL *in vivo* as a non-obligatory holdase. Conspicuously absent from this set are proteins with low pI (<6, polyanions), and class IV proteins, high MW, or many domains (Figure 5A, C, E, G). In all cases, it is because rather than fold slowly, proteins in these categories tend to be intrinsically nonrefoldable. On the other hand, it is interesting to notice an enrichment for very slow refolding (i.e., requiring more than 5 min) for proteins with higher MW (Figure 5E).

With chaperonin, proteins with low pI (<5 or 5–6) are still not particularly slow refolding, but now for the opposite reason: because GroEL is unusually expeditious at refolding them, so they have mostly refolded within 1 min (Figure 5B, Data S2K). Proteins with high pI (>10) show similar kinetics with chaperonin as they do without. This may be because such proteins could bind too tightly to GroEL’s negatively charged lumen, which would render it a less efficient chaperone for these clients (and in a few rare cases, preclude folding). Both class III^-^ and class IV proteins are refolded rapidly by GroEL (Figure 5D), consistent with kinetic models that suggest these proteins form intermediates that rapidly sort to GroEL (Powers et al., 2012; Santra et al., 2017). We find few differences in the rate for folding high-MW or low-MW proteins, a contrast with chaperone-null conditions in which high-MW proteins that fail to refold quickly generally do not recover within 5 min (Figure 5E-F). Finally, with chaperonin, proteins with many domains have more examples that refold slowly, suggesting that GroEL *can* resolve misfolded forms populated by these proteins but likely at the cost of additional annealing cycles (Figure 5H). To summarize, GroEL likely requires a greater metabolic cost and more annealing cycles to refold proteins with many domains and with positive charge.

### GroEL/ES is Crucial for Folding α/β Folds

Because our PK susceptibility measurements can be resolved down to individual residue locations, it is possible to assign nonrefolding sites to specific structural domains within proteins. Using the SCOP database (structural classification of proteins (Gough et al., 2001; Pandurangan et al., 2019)), such domains can be grouped into fold-types, reflecting deep evolutionary relationships between polypeptides that share a common topology despite having very different sequences and functions. The intrinsic refoldability levels of different folds in cyto-serum largely preserve trends previously observed (Figure 5I) (To et al., 2021). In particular, small domains with ‘simple’ topologies (low contact order (Plaxco et al., 1998)) tend to be the most refoldable, such as OB-folds (79%), 3-helical bundles (84%), ubiquitin-like folds (88%), and SH3 barrels (100%). The specialized folds that are unique to aminoacyl-tRNA synthetases (aaRSs) are generally the *least* intrinsically refoldable, namely the adenine nucleotide α-hydrolase-like fold (46%, the core of class I synthetases), and the class II aaRS core fold (21%). TIM barrels display slightly lower-than-average levels of refoldability in cyto-serum (62%, average is 64%).

GroEL has a profoundly restorative effective on these fold-types (Figure 4I), elevating the refolding frequencies of the class I and class II aaRS folds to 83% and 77% respectively. In our experiment, GroEL rescued many TIM barrels (raising their refolding frequency to 81%) which is consistent with the previous observation that GroEL has a strong preference to co-precipitate TIM barrel-containing proteins (Kerner et al., 2005; Georgescauld et al., 2014).

However, we found additionally that GroEL had very pronounced effects on assisting Rossmann-folds (of both the NADH-binding (55% to 87%) and SAM-binding (73% to 100%) sub-lineages), P-loop NTPases (64% to 95%), and PRTase-like domains (29% to 100%). All the fold-types that disproportionately benefit from GroEL have α/β architectures (Cheng et al., 2014; Schaeffer et al., 2017) (with the exception of the class II aaRS fold, which is α+β). In the presence of GroEL, we find that all fold-types are highly refoldable, implying that GroEL smooths over the intrinsic differences in refoldability associated with different protein topologies.

### DnaK is Also a Versatile Chaperone that Complements GroEL

Along with GroEL/GroES, the other key chaperone in *E. coli* is DnaK (Hsp70), which operates with its co-chaperone DnaJ (Hsp40) and a nucleotide exchange factor, GrpE (Rosenzweig et al., 2019; Mayer and Gierasch, 2019). In experiments conceptually similar to those described in the previous sections (Figure 6), we performed global refolding assays in which 5 µM DnaK, 1 µM DnaJ, and 1 µM GrpE were supplemented into the cyto-serum refolding dilution buffer (as well as to the native samples, as in Figure 1A). Initial analysis provided poor coverage (759 proteins total; Figure 3–figure supplement 2A-B), owing to the fact that DnaK, DnaJ, and GrpE (abbreviated as DnaK/J/E) are cleaved by Proteinase K at many locations, and accounted for 1038 (11%) of all peptides quantified. To rectify this matter, we performed a 12-way LFQ in which raw spectra from the three biological replicates of cyto-serum/GroEL+GroES native and refolded, and the three biological replicates of cyto-serum/DnaK+DnaJ+GrpE native and refolded were analyzed together, and peptides identified and quantified in the 6 DnaK channels were then extracted (see Methods).

**Figure 6.**
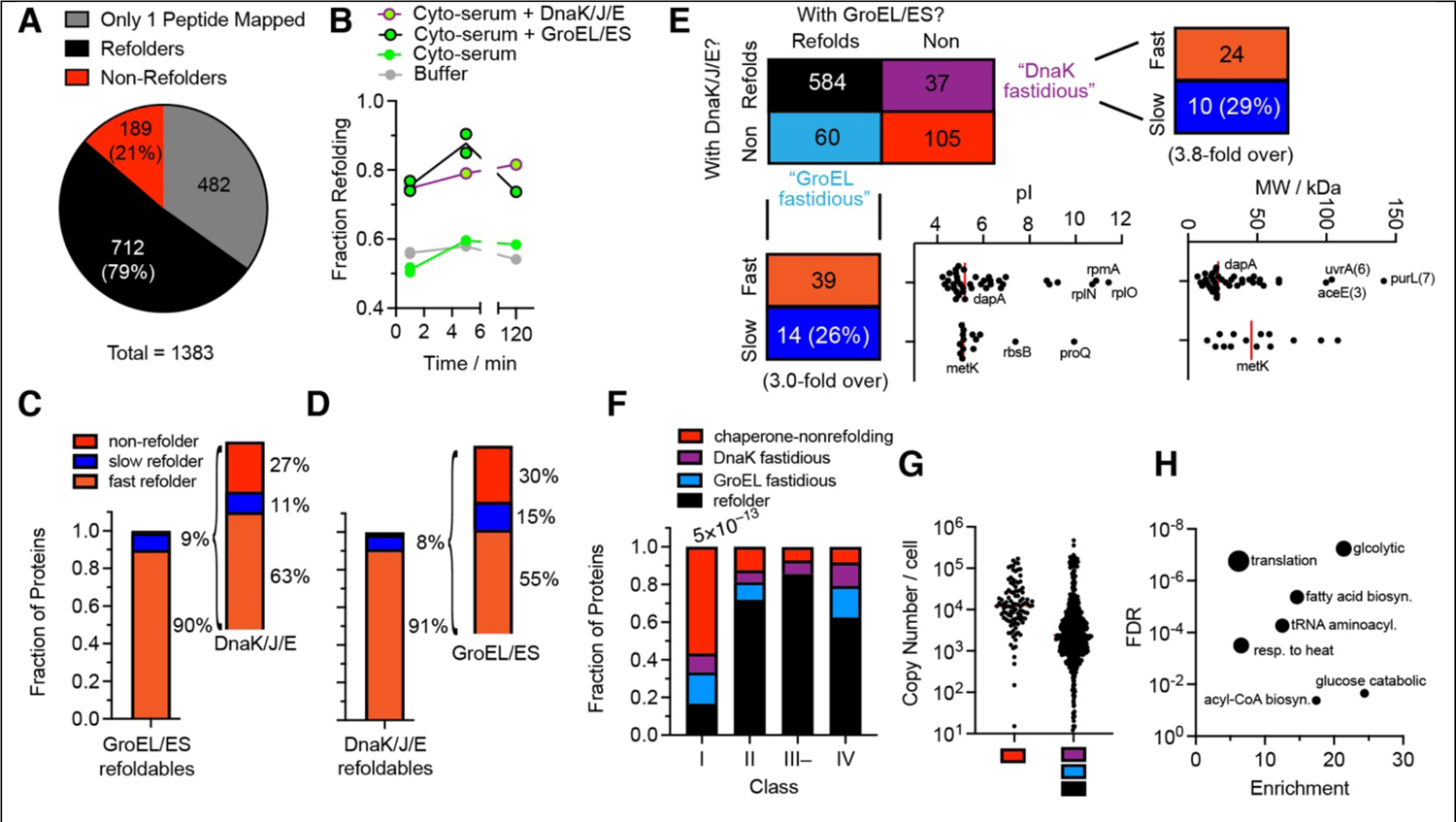
DnaK/DnaJ/GrpE (Hsp70/40) refolds many *E. coli* proteins, with only a few that are fastidious for one chaperone over the other. (A) Pie charts showing the total number of proteins assessed, of which how many are designated refoldable (black; 0 or 1 peptides with significant difference in proteolytic susceptibility in refolded form) or non-refoldable (red; 2 or more peptides with significant difference in proteolytic susceptibility in refolded form) for refolding experiments in cyto-serum with DnaK/J/E. In gray are proteins with only 1 peptide quantified, which are not used in further analysis (or counted in the refolding percentages). Data correspond to #e in Figure 1–figure supplement 1C (Data S3). (B) Fraction of proteins that refold after 1 min, 5 min, or 120 min in either buffer (gray), cyto-serum (green), cyto-serum with GroEL/ES (green, black borders), or cyto-serum with DnaK/J/E (green, purple borders). (C) Frequency of slow refolding with GroEL/ES. This covers the 740 proteins that refolded within 1 min in the cyto-serum/GroEL/ES experiment (called ‘fast refolders’) or did not refold within 1 min but did refold within 5 min in the cyto-serum/GroEL/ES experiment (called ‘slow refolders’). Of the 66 proteins that refold slowly with GroEL, the bar to the right shows the frequency of proteins that refolded fast (within 1 min), slow (not within 1 min but within 5 min), or not at all in the cyto-serum/DnaK/J/E experiments. Data correspond to #4 and #5 in Figure 1–figure supplement 1B. (D) Frequency of slow refolding with DnaK/J/E. This covers the 638 proteins that refolded within 1 min in the cyto-serum/DnaK/J/E experiment (fast refolders), or did not refold within 1 min but did refold within 5 min in the cyto-serum/DnaK/J/E experiment (slow refolders). Of the 49 proteins that refold slowly with DnaK, the bar to the right shows the frequency of proteins that refolded fast, slow, or not at all in the cyto-serum/GroEL experiments. Data correspond to #e and #f in Figure 1–figure supplement 1C. (E) Truth table summarizing the results of the 12-way LFQ pooling 3 replicates of native/cyto-serum/GroEL/ES, 3 replicates of refolded[5min]/cyto-serum/GroEL/ES, 3 replicates of native/cyto-serum/DnaK/J/E, and 3 replicates of refolded[5min]/cyto-serum/DnaK/J/E (Data correspond to #e in Figure 1–figure supplement 1C). Analysis covers 786 proteins for which at least 2 peptides could be confidently quantified in both conditions. Proteins that refold only with GroEL/ES are called “GroEL fastidious” (light blue) and those only with DnaK/J/E are called “DnaK fastidious” (purple). Shown also is, among the DnaK fastidious proteins, how many refold fast or slow with DnaK; and among the GroEL fastidious proteins, how many refold fast or slow with GroEL. pI and MW distributions for the GroEL fastidious proteins are given, broken down by whether they are fast GroEL refolders or slow GroEL refolders. (F) Frequency of proteins that refolded in both conditions (black), only with GroEL/ES (light blue), only with DnaK/J/E (purple), or did not refold in either (chaperone-nonrefolder; red), separated on the basis of chaperonin class (Kerner et al., 2005; Fujiwara et al., 2010). Numbers listed above bars indicate P-value by the chi-square test. (G) Abundance of the 105 chaperone-nonrefolding proteins, compared to the other 681 in this analysis, according to Li et al. (2014). (H) Gene ontology enrichment analysis of the 105 chaperone-nonrefolding proteins, compared to the *E. coli* genome, using PantherDB (Mi et al., 2019)

Through this approach, peptides present in the DnaK/J/E refolding samples that failed to be identified can be identified if they match a feature (in retention time and m/z) that was sequenced in the corresponding GroEL samples. It is important to point out that since our analysis is by LFQ and all samples are injected separately, pseudo-SILAC quantification of DnaK/J/E samples is unaffected by the GroEL samples. With this change, the DnaK experiment’s coverage improved: we could quantify 11445 peptides (Figure 3–figure supplement 2D), making refoldability assessments on 901 proteins (Figure 3–figure supplement 2C), comparable to that of the GroEL experiment (998 proteins, 12562 peptides).

DnaK results in 79% of the *E. coli* proteome refolding after 5 min (Figure 6A, Data S3), comparable but slightly less to that of GroEL (85%). Indeed, virtually all of the refoldability trends we found for GroEL were echoed with DnaK. This includes: a flattened pI-dependence (Figure 6–figure supplement 1A), a flattened MW-dependence with a less pronounced dip at 60-80 kDa (Figure 6–figure supplement 1B), and very little dependence on subunit count (Figure 6–figure supplement 1C). The most salient difference is DnaK is somewhat worse at refolding large >80 kDa proteins (77%) compared to GroEL (91%). Class I proteins remain challenging candidates for DnaK (25% refoldable) and class IV proteins appear to partially benefit from DnaK (refolding at 75%), though maintain a preference for GroEL (which restores 91% of them), as expected (Figure 6–figure supplement 1D).

One feature that is distinct and also quite telling about DnaK is that its time dependence is very different from GroEL’s (Figure 6B). At 1 min, its refolding performance on the *E. coli* proteome is similar to that of GroEL’s, and it proceeds to steadily increase up to 2 h. This is unlike GroEL’s dependence which shows a major increase at 5 min, and then decreases significantly at 2 h. The latter observation could potentially be explained by pointing out that after 2 h, GroEL exhausts its ATP supply whereas DnaK (which uses 1 ATP per cycle rather than 7) might not.

Our results suggest that a refolding problem that is ‘challenging’ for one chaperone is not necessarily challenging for another. For instance, when we look at the minority of GroEL/ES refolders that required more than 1 min to refold (slow refolders, 66 proteins in total), the majority are refolded quickly by DnaK (Figure 6C). Ipso facto, for the minority of DnaK/J/E refolders that required more than 1 min to refold (51 proteins in total), the majority are refolded quickly by GroEL (Figure 6D, Data S3K). Hence, the strengths of these chaperones are complementary for certain clients.

In our 12-way LFQ that includes both the GroEL and DnaK refolding conditions at the 5 min timepoint, we identify 786 proteins for which 2 or more peptides were detected in each condition (Figure 6E), thereby permitting an independent of assessment of refoldability under both conditions (Data SB, see Figure 3–figure supplement 2F for other timepoints). We find that most proteins that refold under GroEL also refold under DnaK, with only a small subset of proteins that appear to be specialized for GroEL (60 total) or DnaK (37 total). We will refer to the clients that can only refold with one chaperone or the other as ‘fastidious’ clients.

Whilst the GroEL-fastidious clients mostly refold rapidly with GroEL (74%), we do find a surprisingly large number that refold slowly with GroEL (26%), 3-fold more frequent than slow GroEL-refolding in general (cf. Figure 6E). It is instructive to divide the GroEL-fastidious clients into subgroups that refold quickly with GroEL and slowly with GroEL. The fast-refolding GroEL-fastidious clients are disproportionately acidic (the median pI of this group is 5.13 with 3 ribosomal proteins discounted) and low-MW (with 3 exceptions, though these high-MW proteins have many smaller domains). These proteins therefore most likely require GroEL’s *foldase* activity (folding inside the cage (Brinker et al., 2001)). On the other hand, those GroEL-fastidious clients that refold slowly are perhaps those with highly entrenched misfolded states that require higher energy inputs to unfold and many iterative annealing cycles to fully correct. These proteins therefore most likely employ GroEL’s *stronger unfoldase* activity. This hypothesis is supported by the fact that this group includes the well-known obligate GroEL client, MetK.

DnaK-fastidious clients also have a surprisingly large number of cases that refold slowly with DnaK (29%), 3.8-fold more frequent than slow DnaK refolding in general. One hypothesis is that these proteins may form misfolded states that aggregate very rapidly, and therefore rely on DnaK’s disaggregase activity, a function that GroEL lacks. Supporting the theory that DnaK can disaggregate proteins in our refolding experiments is the finding that it almost entirely prevents ‘fold-losing’ at later time points (Figure 3–figure supplement 2E, a feature unique to it), and that, where the data was available for comparison, all the DnaK-fastidious proteins (HemB, PyrC, Prs, SerC, Tgt, Ugd) were found to be aggregation-prone in the solubility assays of Niwa et al. (2012).

### Chaperone-Nonrefolders

The most obvious feature of the DnaK/GroEL cross-correlation dataset (Figure 6E) is that there are many proteins that do not refold with *either* GroEL or DnaK, and in fact the most predictive descriptor for whether a protein cannot refold with GroEL is whether it cannot refold with DnaK and ipso facto (odd’s ratio = 51.4; P-value < 10^-66^ by Fisher’s exact test). We refer to this cohort of 105 proteins as ‘chaperone-nonrefolders’ (Data SB). It should be pointed out that some of these proteins could potentially be refolded by other chaperones (Hsp90 and trigger factor) or require multiple chaperones in combination (Langer et al., 1992). We refer to them in the following as chaperone-nonrefolders for brevity’s sake, though what is implied is ‘GroEL/DnaK-nonrefolder.’

Class I proteins are highly over-represented in the cohort of chaperone-nonrefolders (Figure 6F): 59% of class I proteins are chaperone-nonrefolders compared to 14% in general (4.2-fold enriched, P-value < 10^-13^ by chi-square test). Class I proteins are those which were found to be de-enriched from the fraction of proteins that co-precipitate with GroEL/ES (Kerner et al., 2005). Because they do not associate strongly with GroEL *in vivo*, these proteins have historically been construed as efficient intrinsic refolders (Houry et al., 1999; Kerner et al., 2005; Santra et al., 2017). Our data suggests an alternative: that these proteins do not get entrapped within the GroEL/ES cavity because they fold efficiently on the ribosome co-translationally. In so doing, these proteins bypass GroEL (which is preponderantly a post-translational chaperone (Balchin et al., 2005; Houry et al., 1999)). On the other hand, if these proteins became habituated to folding co-translationally, it would also explain why they are recalcitrant to refolding from a full-length denatured form. Whilst there are only a few detailed dissections of obligatory co-translational folding, our data for the five-domain protein EF-G (FusA) agree with a single-molecule study (Liu et al., 2019) in which it was found that folding of domain 1 is ribosome dependent and folding of domain 2 requires folded domain 1. Consistent with this, domains 1 and 2 of FusA are nonrefoldable during our refolding reactions, both with and without chaperones.

Chaperone-nonrefolders are generally very abundant proteins (Figure 6G), and relative to the *E. coli* proteome, the set is greatly enriched for proteins that are involved in core metabolic processes (Figure 6H), such as glycolysis (21-fold enriched, FDR < 10^-7^), fatty-acid biosynthesis (15-fold enriched, FDR < 10^-5^), and tRNA aminoacylation (13-fold enriched, FDR <10^-4^). We also find that this set of proteins is enriched for ‘specialized’ fold-types that have not diversified as broadly. For instance, we find that 10 of the 19 glycolytic-related enzymes are chaperone-nonrefolders (Figure 6H). Amongst this group are enzymes like phosphofructokinase (PfkA), phosphoglycerate kinase (Pgk), and malate dehydrogenase (Mdh). These enzymes each feature specialized folds (e.g., phosphofructokinase-like, phosphoglycerate kinase-like, and LDH C-terminal domain-like) of which in the *E. coli* proteome we only have data on one example, and in all cases that domain is chaperone-nonrefolding. There is also a great over-representation of aminoacyl-tRNA synthetases in this group (8 in total), including: AspS, PheT, GlyS, LeuS, ProS, GlnS, SerS, ThrS. It is notable that all of these synthetases except LeuS and GlnS are class II synthetases. Previous refolding assays on purified ThrS have shown that no combination of GroEL and DnaK can reactivate it beyond ∼50% (Kerner et al., 2005), consistent with it (and possibly class II synthetases in general) as chaperone-nonrefolding. Hence, we conclude that abundant proteins that perform core metabolic functions in glycolysis and translation are especially likely to be chaperone-nonrefolding.

## DISCUSSION

### Revising the Scope of Obligate GroEL Refolders

Our study revises aspects of the consensus model of which *E. coli* proteins employ GroEL for efficient refolding. The consensus model is strongly influenced by the classic work by Kerner et al. (2005), in which rapid depletion of ATP was used to entrap GroEL clients within the *cis* cavity of the GroEL/ES complex. Pull-down on a His-tagged GroES then resulted in co-precipitation of GroEL interactors, which were identified with mass spectrometry. Proteins that were highly enriched in the GroEL fraction, which were termed class III proteins, were found to be generally low-abundance, between 30-60 kDa, and over-represent TIM barrel folds. By analyzing protein refoldability levels in cyto-serum vs. those in cyto-serum supplemented with GroEL and GroES, we can assess which of the proteins that get entrapped with GroEL also depend on it to refold. Our results concur with the finding that TIM barrels tend to be more GroEL-dependent (Figure 6I). On the other hand, the findings that GroEL is particularly important for refolding high-MW and low-pI proteins (Figure 5) in *E. coli* have not been described. Why were these patterns not previously observed? Thoughtful reflection on what pull-down assays can and can*not* show is instructive in this matter. High-MW proteins cannot be entrapped within the sealed GroEL/ES *cis* cavity, and therefore would be systematically excluded from pull-down assays. Indeed, previous work has highlighted several examples in which GroEL restored the activity of high-MW proteins that cannot fit inside the cavity, particularly aconitase (AcnB, 93 kDa) (Chaudhuri et al., 2009), which our study confirms can refold to a native structure in the presence of GroEL. Our experiment also confirms DNA gyrase (GyrA (97 kDa) and GyrB (90 kDa)) and MetE (85 kDa) can refold in the presence of GroEL, notable given that these proteins have been previously shown to interact with GroEL *in vivo* (Houry et al., 1999), even though they do not efficiently co-precipitate with it. Previous work showing that GroEL can refold high-MW proteins has been explained by positing that the *trans* cavity can also bind misfolded clients (Chaudhuri et al., 2009; Farr et al., 2003). Our results suggest that the *trans* mechanism represents a critical function of GroEL. Whilst *E. coli* does not have many proteins with MW greater than 80 kDa, these observations suggest that GroEL plays a significant role in their biogenesis, echoing the observation that eukaryotic TriC/CCT has been shown to principally operate on large proteins (Yam et al., 2008).

A second key feature that emerges from our set of obligate GroEL-refolders is the outsize role GroEL plays in refolding acidic proteins (pI < 6). The negatively-charged cavity walls of GroEL (Tang et al., 2006; Tang et al., 2008) would be expected to create a ‘repulsive field’ for acidic proteins that could facilitate their compaction, overcoming the inter-residue electrostatic repulsion within a protein chain that would counter its tendency to collapse.

Supporting this view is the further observation that the group of slow GroEL refolders has few proteins with low pI (Figures 5B, 6E). Indeed, the primary work which established the potential foldase activity of the GroEL cavity (Brinker et al., 2001) found that inside the cage, GroEL/ES accelerates productive folding (foldase) of *R. rubrum* RuBisCo but merely prevents aggregation of *B. taurus* rhodanese (holdase). Consistent with our model, *Rr*RuBisCo has a low pI (of 5.6) whilst *Bt*Rhodanese does not (6.9). There is a plausible reason why this key relationship with pI was not detected previously: because GroEL refolds acidic protein expeditiously, they would not accumulate within it to become a large steady-state fraction of GroEL occupancy. Such assertions raise the obvious question: What about cationic proteins? Our study shows that *E. coli* protein with high pI are generally efficient intrinsic refolders, and particularly so in the cytosolic medium (Figure 3F and To et al., 2021). These observations highlight the usefulness of assays which characterizes the *impact* of chaperones on the *structural outcome* of their clients, which provide a complementary view to approaches that measure activity, aggregation, or co-precipitation.

### DnaK’s Activities in Relation to GroEL’s

Hsp70s and the menagerie of co-chaperone J-domain proteins have attracted interest in recent years, due to their importance in several diseases and the discovery that they can dissolve amyloid fibrils (Rosenzweig et al., 2019; Gao et al., 2015). Our approach provides a means to compare DnaK’s activity to GroEL’s proteome-wide under the same conditions.

Overall, DnaK and GroEL refold a similar clientele, with only a small number that are specialized (fastidious) for one or the other. These observations are consistent with the prevailing idea that the proteostasis network is integrated (Santra et al., 2017) rather than compartmentalized, with a large amount of redundancy built in: there may be proteins that *prefer* to use DnaK (because it requires less energy), or *prefer* to use GroEL (because they bind to it more rapidly), but ultimately most clients can use either. This finding is consistent with an emerging view that most chaperones share a common mechanism that can be effective on many clients, namely, unfoldase activity on misfolded states (Figure 7A) (Lin et al., 2008; Balchin et al., 2020; Macošek et al., 2021; Imamoglu et al., 2020), thereby providing those molecules with further opportunities to refold properly (the iterative annealing mechanism (Thirumalai and Lorimer, 2001)).

**Figure 7.**
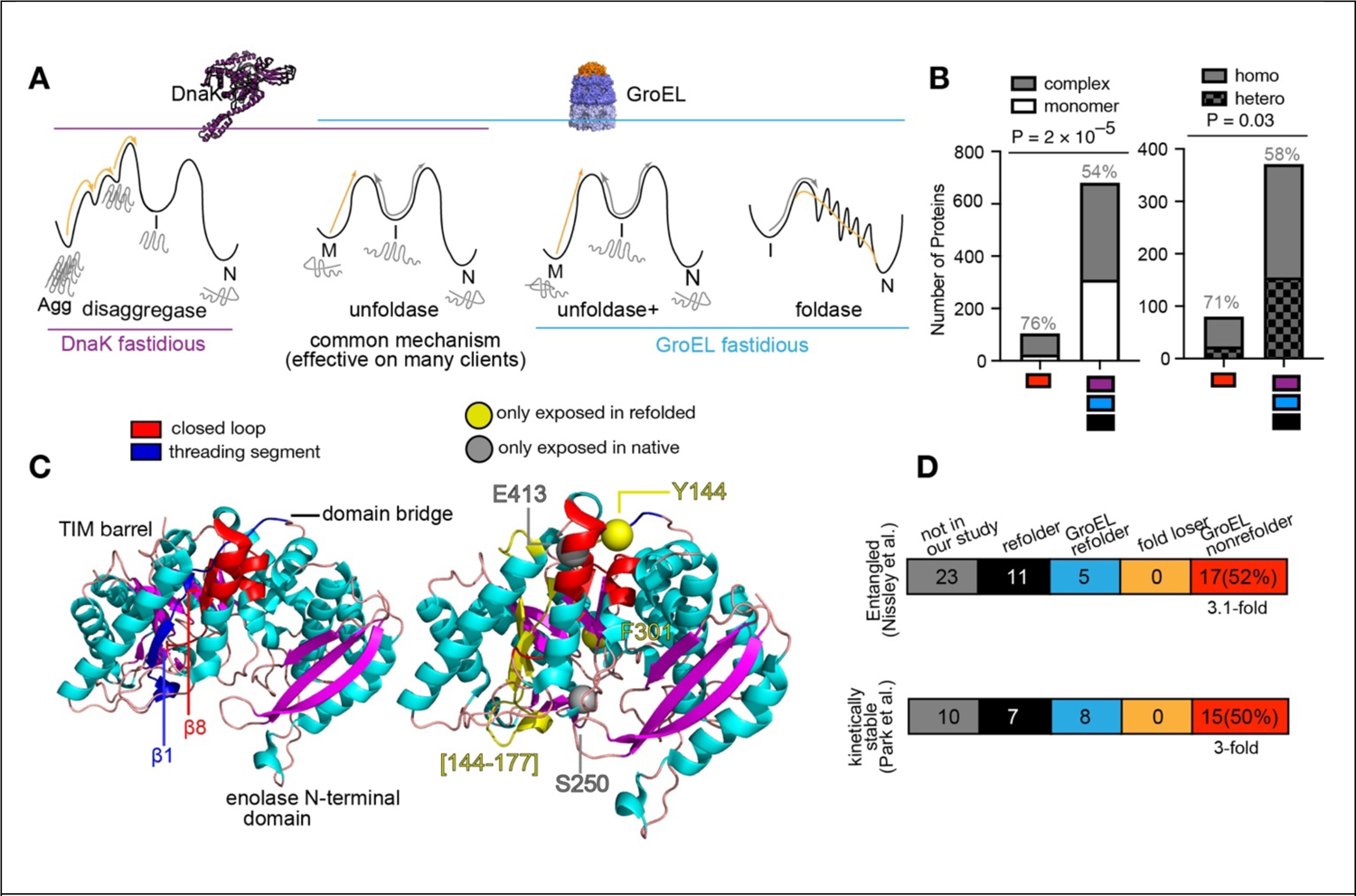
A unified model for chaperone functions in *E. coli*. (A) A model for the overlapping, but distinct, activities of DnaK and GroEL. Four chaperone functions are illustrated along with their effect on a protein’s folding free energy diagram. Chaperone function shown with orange arrows, whereas intrinsic protein behaviour shown with gray arrows. Both DnaK and GroEL can perform unfoldase activity on misfolded states. DnaK is unique for its disaggregase activity. GroEL is unique because its unfoldase activity is stronger, and can resolve more entrenched misfolded states, and because its charged lumen can act as a foldase and catalyze folding, particularly for acidic proteins. (B) Left, Number of proteins that are monomeric or in constitutive complexes for chaperone-nonrefolding proteins (as defined in Figure 5E) or for the other 681 in the analysis. Gray percentages represent fraction in complexes. P-value according to Fisher’s exact test. Right, Number of proteins in complexes that are in homomers or in heteromers for chaperone-nonrefolding proteins or for the other proteins in the analysis. Gray percentages represent fraction in homomers. P-value according to Fisher’s exact test. Data correspond to #e in Figure 1–figure supplement 1C. (C) Left, structure of *E. coli* enolase (PDB: 2FYM) showing the N-terminal domain, bridge, and C-terminal TIM barrel. Computational models predict potential formation of an entangled state whereby β1 (blue) would thread through a loop closed by β8 and the C-terminal α helix (red) (Ritaban et al., 2021). Right, locations of five all-or-nothing peptides identified following refolding for 5 min in cyto-serum with GroEL/ES. Yellow represents regions only susceptible to PK in the refolded form; gray, only susceptible to PK in native form. (D) Further analyses on GroEL-nonrefolding proteins (based on #b in Figure 1–figure supplement 1C), correlating with separate studies which identified proteins that were found in a computational model to form entangled near-native states that would bypass recognition from chaperones (Top; Nissley et al., 2021); or that were found to be kinetically stable by remaining undigested by proteases for days (Bottom; Park et al., 2007). GroEL-nonrefolders are 3.1-fold enriched in the cohort of entangling proteins and 3-fold enriched in the cohort of kinetically stable proteins.

However, DnaK and GroEL also have aspects that make them unique (Figure 7A). In addition to being an unfoldase, DnaK is a disaggregase, whilst GroEL’s cavity can also be a foldase (Balchin et al., 2020; Tang et al., 2006). Moreover, GroEL is a stronger unfoldase because its apical domain movements (which couple to unfolding) are driven by cooperative binding/hydrolysis of 7 ATPs. In some cases, GroEL’s stronger unfoldase function may be required for a handful of clients that populate misfolded states that are deeply energetically-entrenched (with MetK and DapA as important examples).

DnaK is also a disaggregase, a critical function that was probably rendered less important in our assay because of the low aggregation levels we encounter thanks to low concentrations and high complexity in our refolding reactions (Figure 1–figure supplement 2D, 3). It is likely that aggregation occurs less in these complex mixtures compared to experiments with purified proteins because aggregation is more efficient between molecules of the same composition (Vecchi et al., 2020; Bianco et al., 2019). The few DnaK-fastidious refolders are likely those whose misfolded states aggregate rapidly enough (on the min-timescale) that GroEL does not have enough time to intercept them.

### Chaperonins Potentiated A Great Expansion of α/β Folds

Are certain types of protein topologies better at folding themselves than others? Our study suggests that under cellular-like conditions, small all-β domains refold the best, specifically, ubiquitin-like folds, SH3 barrels, and OB-folds. These findings support the theory that all-β domains were the earliest globular proteins, the immediate descendants of amyloids (Petrov et al., 2015; Bowman et al., 2020). On the other hand, the most expansive and versatile folds are all α/β, and include TIM barrels, Rossmanns, and P-loop NTP hydrolases, though these folds all display stronger dependence on GroEL (Cheng et al., 2014). The current view is that Hsp60s (relatives of GroEL) are very ancient, and possibly the only chaperone system the last universal common ancestor (LUCA) possessed (Rebeaud et al., 2021). In light of this, we theorize that these fold-types co-emerged with chaperonin, and that the emergence of chaperonin led to a great expansion of protein functional space attendant with them (Lindquist, 2009). Once these larger, more topologically complex domains could be efficiently folded, their functional versatility became accessible, and they became the most dominant architectures of the protein world. Our study therefore suggests a potential chronology for early protein evolution. Smaller all-β architectures likely preceded the expansion of α/β architectures, which were enabled by the emergence of chaperonins and translation. Aminoacyl-tRNA synthetases are often considered as among the most ancient proteins, but we hypothesize that they represent later additions that emerged in tandem with translation, as their folding appears to be more translation-dependent (Fried et al., 2021). The prevalence of chaperone-nonrefoldability among translation and glycolytic proteins, which played formative roles in early life, alludes to the possibility that these key proteins were products of an early functional translation apparatus.

### Why Are Some Proteins Not Refoldable Even with Chaperones?

The observation that a few *E.* coli proteins cannot fully refold from a denatured state with either GroEL or with DnaK invites conversation about how these proteins locate their native states in the first place. Some possibilities include these proteins require HtpG (*E. coli*’s Hsp90), trigger factor (TF), a combination of DnaK and GroEL, or the full complement of all *E. coli* chaperones. Whilst these explanations cannot be excluded, we favour a more parsimonious explanation: that these chaperone-nonrefolding proteins have a strong preference to fold cotranslationally on the ribosome. Several lines of evidence support this view.

The majority of these proteins are in complexes (80 out of 105, 76%), of which the majority (57 out of 80, 71%) are in homocomplexes (Figure 7B). Homomers have been shown to be the most likely to assemble during translation in a “co-co” fashion (wherein nascent chains assemble whilst both are in translation) (Bertolini et al., 2021). Our study suggests that this mode of assembly may be obligatory in some situations.

Two- and three-domain proteins are not especially represented in this group, suggesting that improper inter-domain compaction can normally be resolved post-translationally with chaperones (Imamoglu et al., 2020; Frydman et al., 1999). On the other hand, proteins with 4 or >5 domains are greatly over-represented in this group (Figure 6–figure supplement 1F), suggesting that for proteins with high domain counts, the vectorial synthesis of translation to decouple domain folding and preclude inter-domain contacts becomes more obligatory (Liu et al., 2019; Han et al., 2007).

The possibility of ‘obligate’ cotranslational folders invites the question as to what makes these proteins challenging for chaperones to rescue them. One potential explanation is that these proteins can populate soluble misfolded states that are ‘native-like,’ evade detection by chaperones and are very slow to resolve on their own. A second possibility is that chaperones can identify these states but ultimately cannot repair them, targeting them for degradation. The first scenario has recently been investigated by O’Brien and co-workers (Halder et al., 2021), who found computationally that *E. coli* isochorismate synthase (EntC), enolase (Eno), Galactitol-1-phosphate dehydrogenase (GatD), MetK, and purine nucleoside phosphorylase (DeoD) are prone to form near-native entangled states that bypass GroEL. In agreement, our study identifies Eno and DeoD as chaperone nonrefolders (as for the others: EntC had too low coverage for inclusion; MetK is GroEL-fastidious; GatD was not detected in intrinsic refolding experiments but fully refolded in all chaperone experiments, supporting previous work showing its misfolded states rapidly aggregate (Kerner et al., 2005)).

The authors of the computational study predicted that upon folding, a subpopulation of enolase is prone to becoming entangled (Figure 7C, Left), specifically by threading a segment that comprises the interdomain bridge and the first β-strand of the TIM barrel (residues 141– 158, blue) between a closed loop formed by the final β-strand of the TIM barrel (382–396, red) and the C-terminal helical region (408–416, red). Focusing on the five all-or-nothing peptides detected for this protein in the GroEL/ES-refolding experiment, we find striking agreement with this prediction. Y144 (adjacent to the crossover point) only becomes susceptible to PK in the refolded state, whereas E413 on the surface of the protein is only susceptible to PK in the native state (Figure 7C, Right). This is consistent with the expected structural transformation attendant the threading region passing through the loop. In the native protein, a long stretch between residues 144–177 (that happens to have no Arg or Lys) is immune to PK, and survives as a full-tryptic fragment. It corresponds to a stable structural region corresponding to an antiparallel β-hairpin within the TIM barrel. This full-tryptic fragment disappears in the refolded sample, consistent with the prediction that it would become exposed to proteolysis upon threading. Two further all-or-nothing half-tryptic peptides are identified, mapping to S250 and F301. These do not appear to be related to the local entanglement discussed here, but they are relatively close to each other and may suggest an additional location in enolase prone to misfold.

In a follow-up study, Nissley et al. (2021) computationally probed a larger cohort of *E. coli* proteins, and identified a group of 57 that are expected to bypass chaperones, not aggregate, and not be degraded on account of potentially populating entangled near-native conformations. Our study of obligatory GroEL refolding (cf. Figure 4) covered 33 of these, of which only 11 were found to be intrinsic refolders (33%), 5 were obligate-GroEL refolders (15%), but 17 were GroEL-nonrefolders (52%, 3.1-fold overrepresented) (Figure 7D). Twelve were found that also could not refold with DnaK (39%, 2.9-fold overrepresented). The sizable enrichment of predicted entangled states in our chaperone-nonrefolding cohort and the structural agreement for enolase provides evidence that entanglement provides some of the structural basis as to why chaperones may not be able to rescue all misfolded proteins.

### Chaperone-Nonrefolders May Have Kinetically-trapped Native States, Obviating the Need for Chaperones After Synthesis

If a protein cannot be refolded, even with chaperones, does it represent a liability to cell? Not necessarily, as long as the protein: (i) folds efficiently for the first time, on the ribosome; and (ii) has an unfolding rate that is so slow to be negligible on the physiological timescale. It makes economic sense why such a constellation of features would be desirable for high-abundance proteins, as obligate chaperone involvement in their biogenesis would carry a greater energetic burden. Our study supports the view that chaperone-nonrefolding proteins might frequently be trapped in their native states, hence explaining why under physiological conditions they would not require the service of chaperones after their translation. Indeed, half of the highly kinetically stable proteins identified by Marqusee and co-workers (that remain undigested by thermolysin or trypsin over several days; Park et al., 2007) are GroEL-nonrefolding in our study (Figure 7D), a 3-fold enrichment. Ten were found to also not refold with DnaK (40%, 3-fold enrichment).

The existence of kinetically-trapped native states that are inefficiently refolded from denatured forms poses a number of interesting consequences for how cells maintain proteostasis. For instance, they could potentially explain why such a large fraction of the proteostasis network is dedicated to degradation rather than restoration. Whilst protein degradation represents, on one hand, an important part of gene regulation; it is also possible that in other situations it responds to a biophysical imperative – of a protein that cannot be repaired and must be resynthesized from scratch, to fold co-translationally anew. The slow decay of kinetically-trapped native states out of their prescribed free energy minima could then encode an intrinsic ‘expiration date,’ effectively defining the desired timescale for which the protein should persist.

### Limitations of this Study

The primary limitation in this study is imperfect coverage. The more peptides we quantify per protein, the more potential we have to elucidate nonrefolding regions, and this coverage is not spread equally over all proteins. Fortunately, peptide-level analysis (Figure 3– figure supplement 3) can assure us that differences in coverage do not bias any of the primary trends we observe with respect to pI, MW, or chaperonin class. It does however explain imperfect reproducibility at the individual-protein level (Figure 3–figure supplement 1). The main reason a protein’s refoldability status would change in a different experimental replicate is that the protein lies close to the cut-off and a given peptide (that had statistically-significantly different abundance in refolded samples) was detected in one replicate of the experiment but not the another. Continuous improvements in proteomics technologies are expected to mitigate this limitation.

## MATERIALS AND METHODS

### Materials

| Reagent of Resource | Source | Identifier |
| --- | --- | --- |
| **Bacterial Strains** |  |  |
| *E. coli* (Strain K12) | New England BioLabs | ER2738 |
| **Chemicals and Reagents** |  |  |
| Adenosine diphosphate (ADP) | Sigma-Aldrich | A2765 |
| Adenosine Triphosphate (ATP)<br>10 mM | New England BioLabs | P0756S |
| Ammonium Bicarbonate | Acros Organics | 393210050 |
| Dithiothreitol (DTT) | Sigma-Aldrich | D06632 |
| Glycerol | Fisher | G33-1 |
| Guanidium Chloride (GdmCl) | Acros Organics | 50-01-1 |
| Hydrochloric Acid (HCl) | Fisher Scientific | A142-212 |
| Iodoacetamide (IAA) | Acros organics | 122271000 |
| KCl | Fisher | BP366-1 |
| MgCl<sub>2</sub> | Fisher | BP214-500 |
| NaCl | Fisher | BP358-212 |
| NAD<sup>+</sup> | Sigma-Aldrich | 481911-5GM |
| Sodium dodecyl sulfate (SDS) | Fisher | BP166-500 |
| Tris-Base | Sigma-Aldrich | RD008 |
| Tris-HCl | Sigma | RD009 |
| Tryptone | Fisher | BP1421-2 |
| Urea | Fisher | U17-212 |
| Yeast Extract | Fisher | BP1422-500 |
| **Components to Make**<br>**Protease Inhibitor Cocktail** |  |  |
| AEBSF.HCl | Thermo Scientific | 78431 |
| Aprotinin from bovine lung | Sigma Aldrich | 10820 |
| Bestatin | Thermo Scientific | 78433 |
| E-64 | Thermo Scientific | 78434 |
| Leupeptin | Sigma Aldrich | E18 |
| Pepstatin A | Thermo Scientific | 78436 |
| **Components to Make MOPS**<br>**MEDIA** |  |  |
| 13C6 L-ARGININE-HCL | Fisher Scientific | PI88433 |
| 13C6 L-LYSINE-2HCL | Fisher Scientific | PI89988 |
| Adenine | Alfa Aesar | A14906 |
| Ammonium Chloride | EMD | AX1270-1 |
| Ammonium Molybdate |  |  |
| Boric Acid | Fisher Scientific | BP-168-1 |
| Calcium Chloride | Fisher | C77-500 |
| Calcium Pantothenate | Acros organics | 24330100000 |
| Cobalt Chloride | Acros | B42357-1000 |
| Cupric Sulfate |  |  |
| Cytosine | TCI America | 71-30-7 |
| di Hydroxy Benzoic Acid |  |  |
| Glucose (Dextrose) (D) | Fisher | D16-3 |
| Guanine | Alfa Aesar | A12024 |
| Iron Sulfate Stock |  |  |
| L-Alanine | Acros organics | 102831000 |
| L-Arginine HCl | Alfa Aesar | A14730 |
| L-Asparagine | Alfa Aesar | b21473 |
| L-Aspartic Acid, Potassium Salt | Alfa Aesar | A13520 |
| L-Cysteine HCl | Alfa Aesar | I06328 |
| L-Glutamic Acid, Potassium Salt | Alfa Aesar | A15031 |
| L-Glutamine | Alfa Aesar | A14201 |
| L-Glycine | J T Baker | 4059-02 |
| L-Histidine HCl H2O | Acros Organics | 166151000 |
| L-Isoleucine | Fisher | bp384-100 |
| L-Leucine | Alfa Aesar | A12311 |
| L-Lysine | Alfa Aesar | J62225 |
| L-Methionine | Alfa Aesar | A10318 |
| L-Phenylalanine | Alfa Aesar | A13238 |
| L-Proline | Fisher | BP-392-100 |
| L-Serine | Acros Organics | 132665000 |
| L-Threonine | Acros Organics | 138931000 |
| L-Tryptophan | Acros Organics | 140590250 |
| L-Tyrosine | Alfa Aesar | A11141 |
| L-Valine | Acros organics | 140811000 |
| Magnesium Chloride | Fisher | BP214-500 |
| Manganese Chloride |  |  |
| MOPS (MW 209.3) | Sigma LS | RDD0018 |
| para-Amino Benzoic Acid | Sigma Aldrich | 1973 |
| para-Hydroxy benzoic Acid | Alfa Aesar | A13700 |
| Potassium Hydroxide | Fisher | P250-3 |
| Potassium Phosphate Dibasic | Fisher | BP363-1 |
| Potassium Sulfate |  |  |
| Sodium Chloride | Fisher | BP358-212 |
| Thiamine | Fisher | BP892-100 |
| Tricine (MW 179.2) | Sigma LS | RDD024 |
| Uracil | Alfa Aesar | A15570 |
| Zinc Sulfate | Alfa Aesar | 33399 |
| **Recombinant Proteins and Enzymes** |  |  |
| DnaK Mix | Cosmo-Biology | GFK-PF003-0.5-EX |
| DNase I from Bovine Pancreas | Sigma Aldrich | 31136 |
| GroEL | Sigma Aldrich | C7688 |
| GroES | Sigma Aldrich | C7438 |
| Endoproteinase LysC | New England BioLabs | P8109S |
| Proteinase K | Fisher Scientific | BP1700 |
| Trypsin -ultra, Mass Spectrometry Grade | New England BioLabs | P8101S |
| **Commercial Assays** |  |  |
| Rapid Gold BCA Assay (Pierce) | Thermo Scientific | A53225 |
| ATP Determination Kit (Invitrogen) | Thermo Scientific | A22066 |
| **LC-MS Reagents** |  |  |
| Acetonitrile (HPLC Grade) | Sigma-Aldrich | 34851 |
| Formic Acid (Optima LC/MS Grade) | Fisher Scientific | A117-50 |
| Trifluoroacetic acid (TFA) (HPLC Grade) | Alfa Aesar | 44630 |
| Water (Optima LC-MS Grade) | Fisher Scientific | W6500 |
| **Databases** |  |  |
| Chaperonin Class Dataset | Kerner et al. 2005 | N/A |
| DnaK Enrichment Dataset | Calloni et al. 2012 | N/A |
| EcoCyc | Keseler et al. 2017 | N/A |
| Fujiwara Data Set | Fujiwara et al. 2010 | N/A |
| SCOP Database | Gough et al., 2001;<br>Pandurangan et al., 2019 | N/A |
| **Instrumentation** |  |  |
| 1/4 Inch Probe | QSonica | N/A |
| 1/8 Inch Probe | QSonica | N/A |
| AN60 Ti Rotor | Beckman Coulter | N/A |
| Eppendorf Centrifuge (5910R) | Eppendorf | N/A |
| OneMP (Mass Photometry) | Refeyn | N/A |
| pH Meter | Mettler-Toledo | N/A |
| Plate Reader (ID3) | Molecular Devices | N/A |
| Plate Reader (Spark) | Tecan | N/A |
| Probe Sonicator (Q700R) | QSonica | N/A |
| Q-Exactive HF-X Orbitrap Mass Spectrometer | Thermo Scientific | N/A |
| Rheometer (MC92) | Anton-Parr | N/A |
| SPEX™ Sample Prep Dual Freezer/Mill | SPEX | N/A |
| SW55 Ti Rotor | Beckman Coulter | N/A |
| UltiMate3000 UHPLC | Thermo Scientific | N/A |
| Ultracentrifuge (Optima XL-A) | Beckman Coulter | N/A |
| Ultracentrifuge (Optima-XPN) | Beckman Coulter | N/A |
| **Other** |  |  |
| 3 mL *Konical Tubes* | Beckman Coulter | N/A |
| 5 mL ultra-clear ultracentrifugation tubes | Beckman Coulter | N/A |
| JL20 Tubes | Beckman Coulter | N/A |
| Mass Spec Vials | Thermo Scientific | N/A |
| Pierce™ 96-Well Polystyrene Plates, White Opaque | Thermo Scientific | 15042 |
| Sep Pak C18 1 cc 50 mg Resin | Waters | 186000308 |
| Viva-Spin 15R 2K MWCO Hydrosart | Sartorius | VS15RH92 |
| 25 mm plate | Anton-Parr | N/A |
| **Software and Algorithms** |  |  |
| AcquireMP | Refeyn | N/A |
| AnalyzerV18 | In House | Github.com/FriedLabJHU/Refoldibility-Tools |
| DCDT+ | John Philo | N/A |
| DiscoverMP | Refeyn | N/A |
| MSFragger Node in PD | Nesvizhskii Lab | N/A |
| Proteome Discoverer v 2.4 (PD) | Thermo Scientific | N/A |
| RheoCompass | Anton-Parr | N/A |
| SEDNTERP | John Philo | N/A |
| XCalibur | Thermo Scientific | N/A |

### RESOURCE AVAILABILITY TABLE

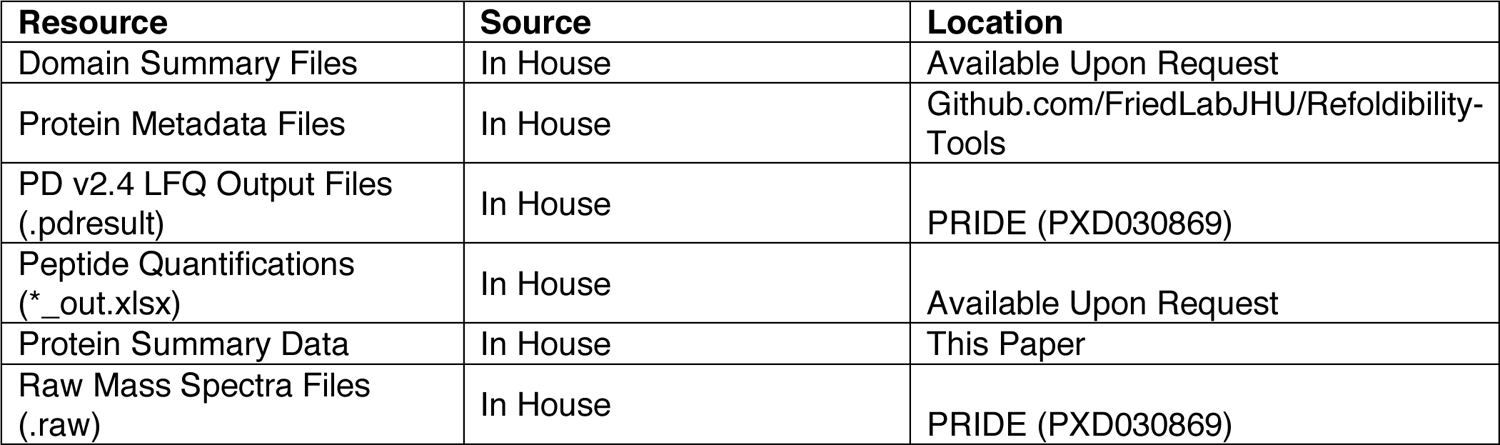

### Methods

#### Preparation of *E.* coli (Strain K12) for Making Cyto-Serum

2 × 1 L of TB Broth (Cold Spring Harbor Recipe) were inoculated with *E. coli* (strain K12) cells in 2 L baffled flask from overnight saturated cultures and grown at 37 °C with agitation (220 rpm) to a final OD_600_ of 2.0. Cells were harvested by centrifugation at 2000 *g* for 15 min at 4°C. Supernatants were removed and cells were washed with 200 mL MPW before centrifuging at 4000 *g* for 15 min at 4 °C. Washed pellets were stored at −20°C until further use.

#### Preparation of K12 for Aggregation Studies

K12 cells were grown in 3 × 50 mL (biological triplicates) of in-house prepared MOPS EZ rich media (lab recipe available upon request; based off of Neidhardt et al., 1977) from saturated overnight cultures with a starting OD_600_ of 0.05. Cells were cultured at 37°C with agitation (220 rpm) to a final OD_600_ of 0.8 before being transferred to 3 × 50 mL falcon tubes and collected by centrifugation at 4000 *g* for 15 mins at 4°C. The supernatants were removed, and cell pellets were stored at −20°C until further use.

#### Preparation of K12 for Limited Proteolysis Mass Spectrometry (LiP-MS) Refolding Studies

K12 cells were grown in 2 sets of 3 × 50 mL (biological triplicates) of in-house prepared MOPS EZ rich media(-Arginine/-Lysine) from saturated overnight cultures with a starting OD_600_ of 0.05. One set was supplemented with 0.5 mM [^13^C_6_]L-Arginine and 0.4 mM [^13^C_6_]L-Lysine and the other with 0.5 mM L-Arginine and 0.4 mM L-Lysine. Cells were cultured at 37°C with agitation (220 rpm) to a final OD_600_ of 0.8. Each heavy/light pair was pooled together and then transferred to 2 × 50 mL falcon tubes and collected by centrifugation at 4000 *g* for 15 mins at 4°C. The supernatants were removed, and cell pellets were stored at −20°C until further use. Further preparation of cyto-serum, aggregation study samples, and refolding experiment samples are described in method details below.

#### General Preparation of Cyto-Serum

Washed *E. coli* pellets were resuspended in MPW to a final volume of 15 mL and lysed by sonication (QSonica Q700R) using a ¼ inch probe for 30 mins on time (pulse: 5 sec on / 5 sec off) at 55% amplitude. Lysed cells were transferred to a JL20 centrifugation tube and clarified at 16000 *g* for 15 min at 4°C. To deplete large macromolecules, cellular lysates were transferred to 5 mL ultra-clear ultracentrifugation tubes and ultracentrifuged at 40000 rpm for 20 h at 4°C without sucrose cushion in a SW-55 Ti rotor. Supernatants were then carefully pipetted from the ultracentrifugation tubes to not disturb the pellet and transferred to 2K MWCO Vivaspin 15R centrifugal filters (Saratorius). The filters were spun at in a swing bucket rotor (Eppendorf 5910R) centrifuge at 3000 *g* for 3 h at 4°C to remove any remaining macromolecules. Standard Tris-Glycine SDS-PAGE (6% stacking, 12% resolving) was used to confirm the removal of all protein molecules that would be stained by Coomassie. The resulting filtrate was concentrated using a Vacufuge Plus (Eppendorf) to a final volume of 1.92 mL, the total volume enclosed within the collective cytoplasms of the original *E. coli* population, creating cyto-serum (2 L of cells at OD_600_ 2.0 comprises of 3.2e12 cells, each cell having 0.6 fL of cytoplasm resulting in 1.92 mL of collected cyto-serum). The cyto-serum is aliquoted, flash frozen in liquid nitrogen and stored at −80°C until further use. The cyto-serum as prepared above is used effectively as a 1.195× stock.

#### Characterization of Cyto-Serum

To determine ATP concentration in cyto-serum, washed *E.* c*oli* pellets were resuspended in MPW to a final volume of 1 mL and lysed by sonication using a 1/8-inch probe for 7 mins on time (pulse: 7 sec on / 14 sec off) at 45% amplitude. Lysed cells were then transferred to a 1.5 mL microfuge tube, clarified 16000 *g* for 15 min at 4°C. The concentration of ATP in cyto-serum was determined using an ATP Determination Assay (Invitrogen) based on the enzymatic activity of luciferase in a microtiter format with a plate reader (Tecan) using known concentrations of ATP ranging from 10 µM to 100 nM as standards. Individual wells of a 96 well opaque polystyrene plate (Thermo Scientific) were loaded with 20 µL of sample and combined with 180 µL of working reagent before incubating at room temperature in the dark for 10 mins prior to luminescence analysis. The amount of ATP in cyto-serum was determined using the obtained concentration of ATP for a 1 mL resuspension volume and dividing by the total volume of cells (50 mL of cells at OD_600_ 2.0 comprises of 8e10 cells, each cell having 0.6 fL of cytoplasm for a total volume of 0.048 mL). The data are reported as a mean ± standard deviations from three independent preparations. Ultraviolet–visible absorption spectra of cyto-serum were obtained with a NanoDrop One (Thermo Scientific) spectrophotometer. The cyto-serum was diluted 60× with MPW prior to analysis and MPW was used as the background.

The pH of cyto-serum was measured for three independent preparations of cyto-serum using a pH meter (Mettler Toledo), using a three-point calibration at pH 4.0, 7.0, and 10.0. The viscosity of cyto-serum was measured via a MRC92 rheometer (Anton-Parr) using 25 mm parallel plates. 25 data points were collected with a point duration of 20 s with a Shear Rate ranging from .01 to 100 1/s logarithmic.

### Preparation of Normalized Lysates

For proteome-wide aggregation and refolding studies, frozen cell pellets were resuspended in a lysis buffer consisting of either 900 µL of Tris pH 8.2 lysis buffer (20 mM Tris pH 8.2, 100 mM NaCl, 2 mM MgCl_2_ and supplemented with DNase I to a final concentration (f.c.) of 0.1 mg mL^-1^) or 900 µL of cyto-serum lysis buffer (1× cyto-serum, supplemented with DNase I to a f.c of 0.1 mg mL^-1^). In samples prepared for analytical ultra-centrifugation as described below, 9 µL of a 100x protease inhibitor cocktail (100 mM AEBSF.HCL, 80 µM Aprotonin, 5 mM Bestatin,1.5 mM E-64.2 mM Leupeptin, and 1 mM Pepstatin A) was added to prevent protein degradation during analysis. Resuspended cells were flash frozen by slow drip over liquid nitrogen and cryogenically pulverized with a freezer mill (SPEX Sample Prep) over 8 cycles consisting of 1 min of grinding (9 Hz), and 1 min of cooling. Pulverized lysates were transferred to 50 mL centrifuge tubes and thawed at room temperature for 20 min. Lysates were then transferred to fresh 1.5 mL microfuge tubes and clarified at 16000 *g* for 15 min at 4 °C to remove insoluble cell debris. To deplete ribosome particles, clarified lysates were transferred to 3 mL *konical* tubes and ultracentrifuged at 33,300 rpm at 4 °C for 90 min without sucrose cushions using a SW55 Ti rotor. Protein concentrations of clarified lysates were determined using the bicinchoninic acid assay (Rapid Gold BCA Assay, Pierce) in a microtiter format with a plate reader (Molecular Devices iD3) using BSA as a calibration standard. Due to the reducing nature of cyto-serum, the BCA assay is incompatible with it. Hence, to determine protein concentrations in lysates prepared in cyto-serum, cell pellets would be generated from the same original liquid culture but split into two equally sized aliquots. The aliquots were resuspended in equal volumes of lysis buffer, with one of the aliquots lysed in Tris native buffer. The two parallel lysates are simultaneously clarified and ultracentrifuged together.

Hence, under these conditions the protein concentration in the Tris-lysed lysate can be used as a surrogate to ascertain protein concentrations in cyto-serum. Generally, the raw the concentrations would be between 3.5 – 4.0 mg mL^-1^ for various preparations. Protein concentrations were diluted to a standard concentration of 3.3 mg mL^-1^ using their respective lysis buffers. This generates the normalized lysates for all downstream workflows.

### Methods to Study Aggregation Preparation of Cell Lysates

For soluble and insoluble protein aggregation studies, native samples were prepared as followed: normalized lysates were diluted with their respective native dilution buffers (20 mM Tris pH 8.2, 100 mM NaCl, 2 mM MgCl_2_, 1.04 mM dithiothreitol (DTT), 62 mM GdmCl; or 1x cyto-serum, 0.1036 mM DTT, 62 mM GdmCl) to a protein concentration of 0.115 mg mL^-1^.

Following dilution, the final concentrations are: 20 mM Tris pH 8.2, 100 mM NaCl, 2 mM MgCl_2_, 1 mM DTT and 60 mM GdmCl; or 1x cyto-serum, 0.1 mM DTT and 60 mM GdmCl. Native samples are then incubated overnight at room temperature. The refolding samples were prepared as follows: 600 μL of normalized lysates, 100 mg GdmCl as a solid, and 2.4 μL of a freshly prepared 700 mM DTT stock solution were combined into a fresh 1.5 mL microfuge tube, and solvent was removed using a Vacufuge Plus to a final volume of 170 μL, such that the final concentrations of all components were 11.6 mg mL^-1^ protein, 6 M GdmCl, 70 mM Tris pH 8.2, 350 mM NaCl, 7 mM MgCl_2_, and 10 mM DTT; or 11.6 mg mL^-1^ protein, 6M GdmCl, 3.5x cyto-serum, and 10 mM DTT. These unfolded lysates were incubated overnight in a sealed container at room temperature to complete unfolding prior to refolding.

### Sedimentation Velocity Analytical Ultracentrifugation

To study the presence of smaller soluble aggregates in refolded extracts using analytical ultracentrifugation, native and unfolded lysates in Tris pH 8.2 were prepared as described above. For analytical ultracentrifugation, all studies were carried out using Tris pH 8.2 refolding buffers as cyto-serum has too many components that absorb at similar wavelengths to proteins (Figure 1–figure supplement 2). To prepare refolded samples, unfolded lysates were diluted 100× with refolding dilution buffer (19.5 mM Tris pH 8.2, 97.5 mM NaCl,1.95 mM MgCl2, and 0.91 mM DTT) and incubated for 2 h at room temperature before being loaded into AUC cells assembled with 1.2 mm double-sector epoxy centerpieces and sapphire windows. Prior to starting each sedimentation velocity (SV) experiment, samples were equilibrated at 20 °C for 1 hour in the centrifuge. Each sample was spun at 20 °C using a 4-hole, An-Ti60 rotor and speed of 50000 rpm. Absorbance was monitored at 280 nm, and radial scans were acquired with 0.003 cm radial steps in continuous mode and with zero time interval between scans. All SV experiments were performed using a Beckman XL-A ultracentrifuge (Beckman Coulter). SV data were analyzed using the time derivative method in dcdt+ (Philo, 2006) to obtain normalized g(s*) distributions. Refolding buffer density (ρ = 1.00464 g/mL) and viscosity (η = 1.0200 cP) were calculated in SEDNTERP (Laue et al., 1992) and an average protein partial specific volume (v ̅) of 0.73 ml g-1 was used to describe the heterogenous cell lysates.

### Mass Photometry (MP)

To monitor smaller soluble aggregates in cyto-serum, Mass Photometry (MP) experiments were conducted on a OneMP instrument (Refeyn) at room temperature. Native samples were prepared in their respective native dilution buffers (either Tris or cyto-serum) as described above. Unfolded samples (either Tris or cyto-serum) were prepared as described above. To prepare refolded samples, 2 µL of unfolded extracts were diluted 100× with 198 µL of refolding dilution buffer (19.5 mM Tris pH 8.2, 97.5 mM NaCl,1.95 mM MgCl_2_, and 0.91 mM DTT; or 0.95× cyto-serum) and incubated at room temperature for 2 h to allow proteins to refold. To prepare samples for MP, 10 µL of native samples or refolding reactions (both at 0.115 mg mL^-1^) were rapidly diluted an additional 100× by addition to 990 µL of Tris lysis buffer and immediately transferred to silicone gaskets on microscope coverslips. Acquisition (which takes 2 min) was initiated within 1 min of the additional 100× dilution. To prepare the set up for sample analysis, microscope cover slips were first cleaned by washing with ethanol, isopropanol, and MPW and then dried with N_2_ gas. Cleaned microscope cover slips were then fitted with a silicone gasket. 10 µL of Tris lysis buffer was loaded onto the silicone gasket to focus and sharpen the instrument. 10 µL of sample was gently pipetted into the droplet seated in the gasket without disturbing focus. Recordings were acquired using the AcquireMP (Refeyn) software and mass distributions were calculated utilizing the DiscoverMP (Refeyn) software.

### Quantification of Pelleting Aggregates Upon Refolding

To study the amount of insoluble aggregates that form upon global refolding, native samples were prepared in their respective native dilution buffers (either Tris or cyto-serum) as described above. Unfolded samples (either Tris or cyto-serum) were prepared as described above. To prepare refolded samples, 5 µL of unfolded extracts were diluted 100× with 495 µL of refolding dilution buffer (19.5 mM Tris pH 8.2, 97.5 mM NaCl,1.95 mM MgCl_2_, and 0.91 mM DTT; or 0.95× cyto-serum) and incubated at room temperature for 2 h to allow proteins to refold (or precipitate). 500 µL of native and refolded samples (both at 0.115 mg mL^-1^, final protein concentration) were centrifuged at 16000 *g* for 15 mins at 4°C to collect aggregated proteins. The supernatant was carefully removed by pipetting to not disturb the protein pellet.

The pellets in all samples were washed with 500 µL of Tris lysis buffer to reduce the interference from reducing agents in Tris or cyto-serum refolding buffers with the BCA assay. The washed pellets were then resuspended in 50 µL of 8M urea in MPW and the protein concentrations were quantified with the BCA Assay as described above. The amount of protein in the pellet was determined using the protein concentration and the resuspension volume (50 µL) and converted to fractional precipitation by dividing by the initial amount of protein in the refolding reaction (57.5 µg). The data are reported as a mean ± standard deviations from biological triplicates, which were differentiated at the inoculation stage. Statistical significance between samples refolded in either Tris or cyto-serum were assessed using t-tests with Welch’s correction for unequal population variances as implemented in Prism 9 (Graphpad). The “precipitation” measured for the native samples were treated as the background level of the measurement because they should not possess any precipitated protein.

### Preparation of Native and Refolded Lysates with and without molecular chaperones for Limited Proteolysis Mass Spectrometry

To prepare half-isotopically-labeled native samples for experiments without molecular chaperones, 3.5 µL of normalized lysates derived from pellets in which half of the cells were grown with [^13^C_6_]L-Arginine and [^13^C_6_]L-Lysine during cell culture and half of the cells were grown with natural abundance L-Arginine and L-Lysine during cell culture (and lysed in cyto-serum), were diluted with 96.5 µL of cyto-serum native dilution buffer (1x cyto-serum, 0.1036 mM DTT, 62.17 mM GdmCl) to a final protein concentration of 0.115 mg mL^-1^. Following dilution, the final concentrations are 1x cyto-serum, 0.1 mM DTT and 60 mM GdmCl. To prepare native samples with the addition of molecular chaperones, 3.5 µL of normalized lysates prepared in cyto-serum were diluted with 96.5 µL of cyto-serum native dilution buffer (1x cyto-serum, 0.1036 mM DTT and 62.17 mM GdmCl supplemented with either 5.19 µM DnaK, 1.04 µM DnaJ and 1.04 µM GrpE; or 4.15 µM GroEL and 8.3 µM GroES) to a protein concentration of 0.115 mg mL^-1^. Following dilution, the final concentrations are 1x cyto-serum, 0.1 mM DTT, 60 mM GdmCl and either 5 µM DnaK, 1 µM DnaJ, 1 µM GrpE; or 4 µM GroEL, 8 µM GroES. While preparing both native and refolding dilution buffers, molecular chaperones were added as the final component and used immediately to prevent them from prematurely utilizing all available ATP. Native samples were then equilibrated by incubating for 90 min at room temperature prior to limited proteolysis. We note here as an important detail that because cyto-serum dilution buffers containing chaperones must be used immediately, and because it is important for reproducibility’s sake that the same buffer preparation is used for all samples, these experiments require three experimentalists working simultaneously to process the three biological replicate samples at the same time.

The refolding samples were prepared as follows: 600 μL of normalized lysates, 100 mg of solid GdmCl, and 2.4 μL of a freshly prepared 700 mM DTT stock solution were added to a fresh 1.5 mL microfuge tube, and solvent was removed using a vacufuge plus to a final volume of 170 μL, such that the final concentrations of all components were 11.6 mg mL^-1^, 6M GdmCl, 3.5x cyto-serum, and 10 mM DTT. These unfolded lysates were incubated overnight at room temperature to complete unfolding prior to refolding.

As above, refolding samples were prepared with or without the addition of molecular chaperones. To prepare refolding samples without molecular chaperones, 99 µL of refolding dilution buffer (0.975x cyto-serum) were added to a fresh 1.5 mL microfuge tube. 1 µL of unfolded extract was then added to the tube containing the refolding dilution buffer and quickly mixed by rapid vortexing, diluting the sample by 100x, followed by flash centrifugation to collect liquids to the bottom of the tube. The final concentrations were 1x cyto-serum, 0.1 mM DTT and 60 mM GdmCl. To prepare refolding samples with the addition of molecular chaperones, 99 µL of refolding dilution buffer (0.975x cyto-serum supplemented with either 5.05 µM DnaK, 1.01 µM DnaJ and 1.01 µM GrpE; or 4.04 µM GroEL and 8.08 µM GroES) were added to a fresh 1.5 mL microfuge tube. 1 µL of unfolded lysate was then added to this refolding dilution buffer and quickly mixed by rapid vortexing, diluting the sample by 100x, followed by flash centrifugation to collect liquids to the bottom of the tube. The final concentrations were 1x cyto-serum, 0.1 mM DTT, 60 mM GdmCl and either 5 µM DnaK, 1 µM DnaJ, 1 µM GrpE; or 4 µM GroEL, 8 µM GroES. Refolded samples were then incubated at room temperature for 1min, 5 min or 2 h to allow for proteins to refold prior to limited proteolysis.

### Limited Proteolysis Mass Spectrometry Sample Preparation

To perform limited proteolysis, 2 µL of a PK stock (prepared as a 0.067 mg mL^-1^ PK in a 1:1 mixture of Tris lysis buffer and 20% glycerol, stored at −20°C and thawed at most only once) were added to a fresh 1.5 mL microfuge tube. After refolded proteins were allowed to refold for the specified amount of time (1 min, 5 min, or 2 h), or native proteins were allowed their 90 min equilibration, 100 µL of the native/refolded lysates were added to the PK-containing microfuge tube and quickly mixed by rapid vortexing (enzyme:substrate ratio is a 1:100 w/w ratio (Feng et al., 2014)), followed by flash centrifugation to collect liquids to the bottom of the tube. Samples were incubated for exactly 1 min at room temperature before transferring them to a mineral oil bath preequilibrated at 110°C for 5 min to quench PK activity. Boiled samples were then flash centrifuged (to collect condensation on the sides of the tube), and transferred to fresh 1.5 mL microfuge tube containing 76 mg urea such that the final urea concentration was 8 M and the final volume was 158 µL. They are then vortexed to dissolve the urea to unfold all proteins and quench any further enzyme activity indefinitely, and flash centrifuged to collect liquids to the bottom of the tubes. Addition to urea is the only allowed pause point; all samples operate on a strict timetable from the moment they are refolded until this point. Moreover, once chaperones are added to cyto-serum, they must be used immediately: in the case of native samples, cyto-serum native dilution buffers are added to proteins immediately after preparation, and then 90 min incubation begins. In the case of refolded samples, cyto-serum refolding buffers are added to unfolded proteins immediately after preparation, and then refolding times (1 min, 5 min, 120 min) begins. This method generates all limited proteolysis samples for this study. For the final studies used for the primary datasets, 51 separate samples were prepared for this experiment, they include: native and refolded in cyto-serum with and without molecular chaperones (DnaK/DnaJ/GrpE or GroEL/ES), and the appropriate biological triplicates for each category. In addition, native samples in cyto-serum prepared with and without GroEL/ES were each prepared on two separate occasions, creating a set of technical duplicates. Refolded samples for each of the three refolding timepoints were prepared in biological triplicates. The 1 min refolding timepoint in cyto-serum with and without GroEL/ES were each prepared on two separate occasions, creating a set of technical duplicates. An additional set was prepared for the 5 min refolding in cyto-serum with the addition of GroEL/ES. A representation of all samples prepared for this study is presented in Figure 1–figure supplement 1. We note here that LiP-MS studies typically prepare a series of parallel ‘control’ samples in which PK is withheld; these samples are then used for standard quantitative proteomics experiments to measure protein abundance differences across conditions (Feng et al., 2014; To et al., 2021). We opted to not perform this for the current study for the following reasons: (1) there is no practical way native and refolded samples that are compared to each other can have different protein abundances given that they are derived from the same lysates; indeed, the samples compared to each other for these studies are compositionally identical and differ only in history; (2) our previous study (To et al., 2021) confirmed that refolded/native protein abundance ratios were equal to unity at a frequency higher than the false discovery rate.

All protein samples were prepared for mass spectrometry as follows: 2.25 μL of a freshly prepared 700 mM stock of DTT were added to each sample-containing microfuge tube to a final concentration of 10 mM. Samples were incubated at 37°C for 30 minutes at 700 rpm on a thermomixer to reduce cysteine residues. 9 μL of a freshly prepared 700 mM stock of iodoacetamide (IAA) were then added to a final concentration of 40 mM, and samples were incubated at room temperature in the dark for 45 minutes to alkylate reduced cysteine residues. To assist trypsin in the digestion of samples with the addition of molecular chaperones, 1 µL of 0.4 µg µL^-1^ Lys-C (NEB) stock was added (enzyme:substrate ratio of 1:100 w/w) and digestion proceeded for 2 h at 37°C. After digestion with Lys-C, 471 μL of 100 mM ammonium bicarbonate (pH 8) were added to the samples to dilute the urea to a final concentration of 2 M. 2 μL of a 0.4 µg µL^-1^ stock of Trypsin (NEB) were added to the samples (to a final enzyme:substrate ratio of 1:50 w/w) and incubated overnight (15-16 h) at 25°C at 700 rpm (not 37°C, so as to minimize decomposition of urea and carbamylation of lysines).

### Desalting of Mass Spectrometry Samples

Peptides were desalted with Sep-Pak C18 1 cc Vac Cartridges (Waters) over a vacuum manifold. Tryptic digests were first acidified by addition of 16.6 μL trifluoroacetic acid (TFA, Acros) to a final concentration of 1% (vol/vol). Cartridges were first conditioned (1 mL 80% ACN, 0.5% TFA) and equilibrated (4 x 1 mL 0.5% TFA) before loading the sample slowly under a diminished vacuum (ca. 1 mL/min). The columns were then washed (4 x 1 mL 0.5% TFA), and peptides were eluted by addition of 1 mL elution buffer (80% ACN, 0.5% TFA). During elution, vacuum cartridges were suspended above 15 mL conical tubes, placed in a swing-bucket rotor (Eppendorf 5910R), and spun for 3 min at 350 g. Eluted peptides were transferred from Falcon tubes back into microfuge tubes and dried using a vacuum centrifuge (Eppendorf Vacufuge). Dried peptides were stored at −80°C until analysis. For analysis, samples were vigorously resuspended in 0.1% FA in Optima water (ThermoFisher) to a final concentration of 0.5 mg mL^-1^.

### LC-MS/MS Acquisition

Chromatographic separation of digests were carried out on a Thermo UltiMate3000 UHPLC system with an Acclaim Pepmap RSLC, C18, 75 μm × 25 cm, 2 μm, 100 Å column. Approximately, 1 μg of protein was injected onto the column. Thecolumn temperature was maintained at 40 °C, and the flow rate was set to 0.300 μL min^−1^ for the duration of the run. Solvent A (0.1% FA) and Solvent B (0.1% FA in ACN) were used as the chromatography solvents. The samples were run through the UHPLC System as follows: peptides were allowed to accumulate onto the trap column (Acclaim PepMap 100, C18, 75 μm x 2 cm, 3 μm, 100 Å column) for 10 min (during which the column was held at 2% Solvent B). The peptides were resolved by switching the trap column to be in-line with the separating column, quickly increasing the gradient to 5% B over 5 min and then applying a 95 min linear gradient from 5% B to 25% B. Subsequently, the gradient was increased from 35% B to 40% B over 25 min and then increased again from 40% B to 90% B over 5 min. The column was then cleaned with a sawtooth gradient to purge residual peptides between runs in a sequence.

A Thermo Q-Exactive HF-X Orbitrap mass spectrometer was used to analyze protein digests. A full MS scan in positive ion mode was followed by 20 data-dependent MS scans. The full MS scan was collected using a resolution of 120000 (@ m/z 200), an AGC target of 3E6, a maximum injection time of 64 ms, and a scan range from 350 to 1500 m/z. The data-dependent scans were collected with a resolution of 15000 (@ m/z 200), an AGC target of 1E5, a minimum AGC target of 8E3, a maximum injection time of 55 ms, and an isolation window of 1.4 m/z units. To dissociate precursors prior to their reanalysis by MS2, peptides were subjected to an HCD of 28% normalized collision energies. Fragments with charges of 1, 6, 7, or higher and unassigned were excluded from analysis, and a dynamic exclusion window of 30.0 s was used for the data-dependent scans. For pseudo-SILAC samples, mass tags were enabled with Δm of 2.00671 Th, 3.01007 Th, 4.01342 Th, and 6.02013 Th (to account for the fixed 6 or 12 Da mass shifts in different charge states) to promote selection of non-chaperone-derived peptides for isolation and data-dependent MS2 scans.

### LC-MS/MS Data Analysis

Proteome Discoverer (PD) Software Suite (v2.4, Thermo Fisher) and the Minora Algorithm were used to analyze mass spectra and perform Label Free Quantification (LFQ) of detected peptides. Default settings for all analysis nodes were used except where specified. The data were searched against Escherichia coli (UP000000625, Uniprot) reference proteome database. For peptide identification, either the PD Sequest HT node (for non-pseudo-SILAC samples) or PD MSFragger node (pseudo-SILAC) were used, each using a semi-tryptic search allowing up to 2 missed cleavages. A precursor mass tolerance of 10 ppm was used for the MS1 level, and a fragment ion tolerance was set to 0.02 Da at the MS2 level for both search algorithms. For Sequest HT, a peptide length between 6 and 144 amino acid residues was allowed. For MSFragger, a peptide length between 7 and 50 amino acid residues was allowed with a peptide mass between 500 and 5000 Da. Additionally, a maximum charge state for theoretical fragments was set at 2 for MSFragger. Oxidation of methionine and acetylation of the N-terminus were allowed as dynamic modifications while carbamidomethylation on cysteines was set as a static modification. For pseudo-SILAC samples, heavy isotope labeling (^13^C_6_) of Arginine and Lysine were allowed as dynamic modifications. All parameters for Sequest HT and MSFragger search algorithms are provided in the table below. The Percolator PD node was used for FDR validation for peptides identified with the Sequest HT search algorithm. For peptides identified with the MSFragger search algorithm, the Philosopher PD node was used for FDR validation. Raw normalized extracted ion intensity data for the identified peptides were exported from the .pdResult file using a three-level hierarchy (protein > peptide group > consensus feature). These data were further processed utilizing custom Python analyzer scripts (available on GitHub, and described in depth previously in To et al., 2021). Briefly, normalized ion counts were collected across the refolded replicates and the native replicates for each successfully identified peptide group. Effect sizes are the ratio of averages (reported in log_2_) and P-values (reported as –log_10_) were assessed using *t* tests with Welch’s correction for unequal population variances. Missing data are treated in a special manner. If a feature is not detected in all three native (or refolded) injections and is detected in all three refolded (or native) injections, we use those data, and fill the missing values with 1000 (the ion limit of detection for this mass analyzer); this peptide becomes classified as an all-or-nothing peptide. If a feature is not detected in one out of six injections, the missing value is dropped. Any other permutation of missing data (e.g., missing in two injections) results in the quantification getting discarded. In many situations, our data provide multiple independent sets of quantifications for the same peptide group. This happens most frequently because the peptide is detected in multiple charge states or as a heavy isotopomer. In this case, we calculate effect size and P-value for all features that map to the same peptide group. If the features all agree with each other in sign, they are combined: the quantification associated with the median amongst available features is used and the P-values are combined with Fisher’s method. If the features disagree with each other in sign, the P-value is set to 1. Coefficients of variation (CV) for the peptide abundance in the three replicate refolded samples are also calculated. Analyzer returns a file listing all the peptides that can be confidently quantified, and provides their effect-size, P-value, refolded CV, proteinase K site (if half-tryptic), and associated protein metadata.

|  | **<u>SequestHT</u>** | **<u>MSFragger</u>** |
| --- | --- | --- |
| **Peak Matching and Outputting** |  |  |
| Precursor Mass Tolerance | 10 ppm | 10 ppm |
| Fragment Mass Tolerance | 20 ppm | 20 ppm |
| Use Average Precursor Mass | FALSE | N/A |
| Use Average Fragment Mass | FALSE | N/A |
| Isotope Error | N/A | 0/1/2 |
| Localized Delta Mass | N/A | 0 |
| Delta Mass Exclude Ranges | N/A | (-1.5,3,5) |
| Max. Precursor Charge | N/A | 4 |
| **Digestion** |  |  |
| Enzyme | Trypsin (semi) | Trypsin (semi) |
| Min. Peptide Length | 6 | 7 |
| Max. Peptide Length | 144 | 50 |
| Max. Missed Cleavage Sites | 2 | 2 |
| Min. Peptide Mass | N/A | 500 |
| Max. Peptide Mass | N/A | 5000 |
| Maximum Charge State for Theoretical Fragments to Match | N/A | 2 |
| Clip n Term M | N/A | TRUE |
| **Spectrum Matching** |  |  |
| Use of Neutral Loss a Ions | TRUE | N/A |
| Use of Neutral Loss b Ions | TRUE | N/A |
| Use of Neutral Loss y Ions | TRUE | N/A |
| Use Ranking Ions | TRUE | N/A |
| Weight of a Ions | 0 | N/A |
| Weight of a Ions | 1 | N/A |
| Weight of a Ions | 0 | N/A |
| Weight of a Ions | 0 | N/A |
| Weight of a Ions | 1 | N/A |
| Weight of a Ions | 0 | N/A |
| **Spectral Processing** |  |  |
| Min. Peaks | N/A | 15 |
| Min. Fragments Modeling | N/A | 2 |
| Min. Ratio | N/A | 0.01 |
| Use Top N Peaks | N/A | 150 |
| Min. Matched Fragments | N/A | 3 |
| Min. Clear m/z Range | N/A | 0 |
| Max. Clear m/z Range | N/A | 0 |
| Mass Calibration | N/A | On and find optimal parameters |
| **Modifications** |  |  |
| Max. Equal Modifications Per Peptide | 3 | N/A |
| Multiple Variable Mods on Residue | N/A | TRUE |
| Max. Variable Mods per Peptide | N/A | 3 |
| Max. Variable Mods Combinations | N/A | 5000 |
| Mass Offsets | N/A | 0 |
| **Dynamic Modifications** |  |  |
| Oxidation | +15.995 Da (M) | +15.995 Da (M) |
| Label: 13C6 | N/A | +6.020 Da (K,R) |
| Acetyl | +42.011 Da (Any N-Terminus) | +42.011 Da (Any N-Terminus) |
| **Static Modifications** |  |  |
| Carbamidomethyl | +57.021 Da (C) | +57.021 Da (C) |

### Refoldability Analysis

Results from analyzer are digested in the following way. Proteins with only one peptide confidently quantified are discounted; proteins with more than two are kept. Peptides are considered to have significantly different abundance in the refolded sample if the effect size is 2 or greater (more than double or less than half the abundance of native), and the P-value is less than 0.01 by Welch’s *t* test. All-or-nothing peptides must have abundance differences greater than 64-fold, and use a relaxed P-value cut-off of 0.0158. The number of significant and all-or-nothing peptides is counted for each protein (or, in the case of Figure 6I, for each domain, whose residue ranges are provided and where peptides are only assigned to a given domain if the PK cut site or the full tryptic range falls within the domain boundaries). Proteins (or domains) are deemed nonrefoldable if two or more peptides with significantly different abundances in the refolded sample are identified.

Protein-level refoldability analyses proceed by counting the number of refoldable and nonrefoldable proteins within a set of categories (e.g., 5 < pI < 6) associated with a feature (e.g., pI) and calculating the fraction refolding within the category. To determine if there is a significant enrichment for (non-)refolders within certain categories, we calculate the expected number of (non-)refolders for each category by taking the total number of proteins that are assigned a value under the feature in question, times the fraction (non-)refolding, times the fraction of proteins in that category. The chi-square test is used to determine if the observed counts and expected counts significantly differ, for all cases in which the feature has three or more categories. If it only has two, Fisher’s exact test is used instead.

Peptide-level refoldability analyses are performed in a similar way. The total number of significant and nonsignificant peptides mapped to proteins within a set of categories associated with a feature are counted and the percentage significant calculated. To determine if there is a significant enrichment for (non-)significant peptides associated with certain categories, we calculate the expected number of (non-)significant peptides for each category by taking the total number of peptides associated with proteins that are assigned a value under the feature in question, times the fraction of peptides that are (non-)significant, times the fraction of peptides associated with that category. The chi-square test is used to determine if the observed counts and expected counts significantly differ, for all cases in which the feature has three or more categories. If it only has two, Fisher’s exact test is used instead.

For condition comparisons (i.e., comparing Tris to cyto-serum, or refolding with GroEL/ES vs. DnaK/J/E), we performed 12-way LFQs, and created a slightly modified analyzer script that assesses peptide quantifications separately for the six samples associated with condition 1 and the six samples associated with sample 2. The analyzer returns a file listing all the peptides that can be confidently quantified, and provides their effect-size, P-value and refolded CV for condition 1 and 2, proteinase K site (if half-tryptic), and associated protein metadata.

Similar to before the number of significant and all-or-nothing peptides are counted for each protein in condition 1 and 2. Proteins are only admitted into the comparison if 2 or more peptides are identified in both conditions, and are classified as refolding in both, refolding in condition 1, refolding in condition 2, or nonrefolding in both. Proteins are discarded if they are on the border; e.g., one significant peptide assigned in condition 1 and two significant peptides assigned in condition 2.

For these analyses, we count the number of proteins associated with a given category (e.g., 5 < pI < 6) that refold in both, refold in condition 1, refold in condition 2, or do not refold in either. For each category, expected counts are calculated by taking the total number of proteins in that category times the overall fraction of proteins that refold in both, refold in condition 1, refold in condition 2, or do not refold in either. The chi-square test is used to determine if the observed counts and expected counts significantly differ. Note that these tests are conducted on individual categories (e.g., the 5 < pI < 6 category is enriched for proteins that refold with GroEL/ES but not without it), whereas previously, the test is conducted on the feature overall (e.g., pI groups do not all refold with the same frequency).

For kinetic comparisons (i.e., comparing proteins that have refolded in cyto-serum for 1 min or 5 min), we combined results from the separate timepoints by collecting the subset of proteins that were identified in both experiments and compiling together the number of significant and all-or-nothing peptides that are counted for each protein at timepoint 1 and 2. Proteins are only admitted into the comparison if 2 or more peptides are identified at both timepoints, and are classified as refolding in both (fast refolder), refolding at the later timepoint (slow refolder), or refolding at the earlier timepoint (fold loser). Nonrefolders are not used for kinetic comparisons. Proteins are discarded if they are on the border; e.g., one significant peptide assigned at timepoint 1 and two significant peptides assigned in timepoint 2. The analyses and chi-square tests are done analogously as above, for the condition comparisons.

### Bioinformatics

Ecocyc database (Keseler et al., 2017) was used to obtain information about cellular compartment (cytosol, inner membrane, periplasmic space, outer membrane, ribosome, cell projection), subunit composition, essentiality, copy number, cofactors, and molecular weight (from nucleotide sequence) for each protein. When the information was available, we used Ecocyc’s Component Of category to obtain the full constitutive composition of the protomer within a complex.

Copy number information predominantly comes from a single ribosome profiling study by Li and co-workers (2014). We used copy number in Neidhardt EZ rich defined medium because of its similarity to the growth medium used in these studies.

Domain information was based on the SCOP hierarchy and obtained through the Superfamily database (http://supfam.org) (Gough et al., 2001; Pandurangan et al., 2019). We used custom scripts to edit the “raw” file available from supfam. org into a format more usable for our purposes (including the switch from a Uniprot identifier to the gene symbol identifier). This database was used to count the number of domains per protein, and to perform the domain-level analysis in which peptides are mapped to individual domains within proteins based on residue ranges. Domains are categorized by their ‘fold.’ Note that in SCOP, folds correspond to collections of superfamilies with similar topologies, and in most situations (but not always) correspond to deep evolutionary relationships (Cheng et al., 2014).

Gene ontology analysis was conducted using PantherDB (Mi et al., 2019). The set of 105 chaperone-nonrefolders was entered as the test set, and the *E. coli* proteome used as the reference set. Statistical overrepresentation tests were selected using the complete set of GO biological processes.

Isoelectric effects were obtained from the isoelectric database (Kozlowski, 2017). We downloaded the file corresponding to E. coli K-12 MG1655 and took an average of the isoelectric points calculated by all the algorithms available for each protein. Chaperonin classes were obtained from Kerner et al. (2005). Specifically, we examined Table S3, manually identified the current Uniprot accession code for each of the proteins identified by Kerner et al., and transferred this information into a file that contains the gene symbol, the current Uniprot accession code, and the class assignment. We also compiled information from Fujiwara et al. (2010) which breaks down class III proteins into class III− and class IV.

## QUANTIFICATION AND STATISTICAL ANALYSIS

All analyses of aggregation were conducted on independent refolding reactions from independent biological replicates (n = 3). Raw values shown for pelleting assay and significance by *t* test with Welch’s correction for unequal population variances. Analytical ultracentrifugation and mass photometry data shown from representative examples from among replicates.

Standard target-decoy based approaches were used to filter protein identifcations to an FDR < 1%, as implemented by Percolator (when searching with Sequest), or Philosopher (when searching with MSFragger).

All mass spectrometry experiments were conducted on three biological replicates used to generate three native samples and three independent refolding reactions from the same biological replicates. For each peptide group, abundance difference in refolded relative to native was judged by the *t* test with Welch’s correction for unequal population variances. Fisher’s method was used to combine P-values when there were multiple quantifiable features per peptide group. P-values less than 0.01 were used as a requirement to consider a region structurally distinct in the refolded form. Differences in means of distributions are assessed with the Mann-Whitney rank-sum test. To test whether particular categories are enriched with (or de-enriched with) (non)refoldable proteins, the chi-square test or Fisher’s exact test is used.

## ACKNOWLEDGMENTS

We thank Susan Marqusee, Ed O’Brien, and Dan Nissley for thoughtful discussion. We thank Philip Mortimer for maintaining the Mass Spectrometer Facility at JHU Department of Chemistry. We thank Di Wu and Grzegorz Piszczek at the National Institutes of Health (Bethesda, MD) for expertise on and assistance with mass photometry experiments. S.D.F. acknowledges support from the NIH Director’s New Innovator Award (DP2GM140926) and from the NSF Division of Molecular and Cellular Biology (MCB2045844). K.G.F. acknowledges support from NIGMS (R01GM079440). T.D. was supported by an NIH training grant (T32GM008403).

## AUTHOR CONTRIBUTIONS

S.D.F. designed the study. P.T., Y.X., T.D., and S.D.F. performed the experiments. All authors analyzed data. P.T. and S.D.F. prepared figures. K.G.F. and S.D.F. secured funding. P.T. and S.D.F. wrote the paper.

## FIGURE SUPPLEMENTS

**Figure 1– figure supplement 1.**
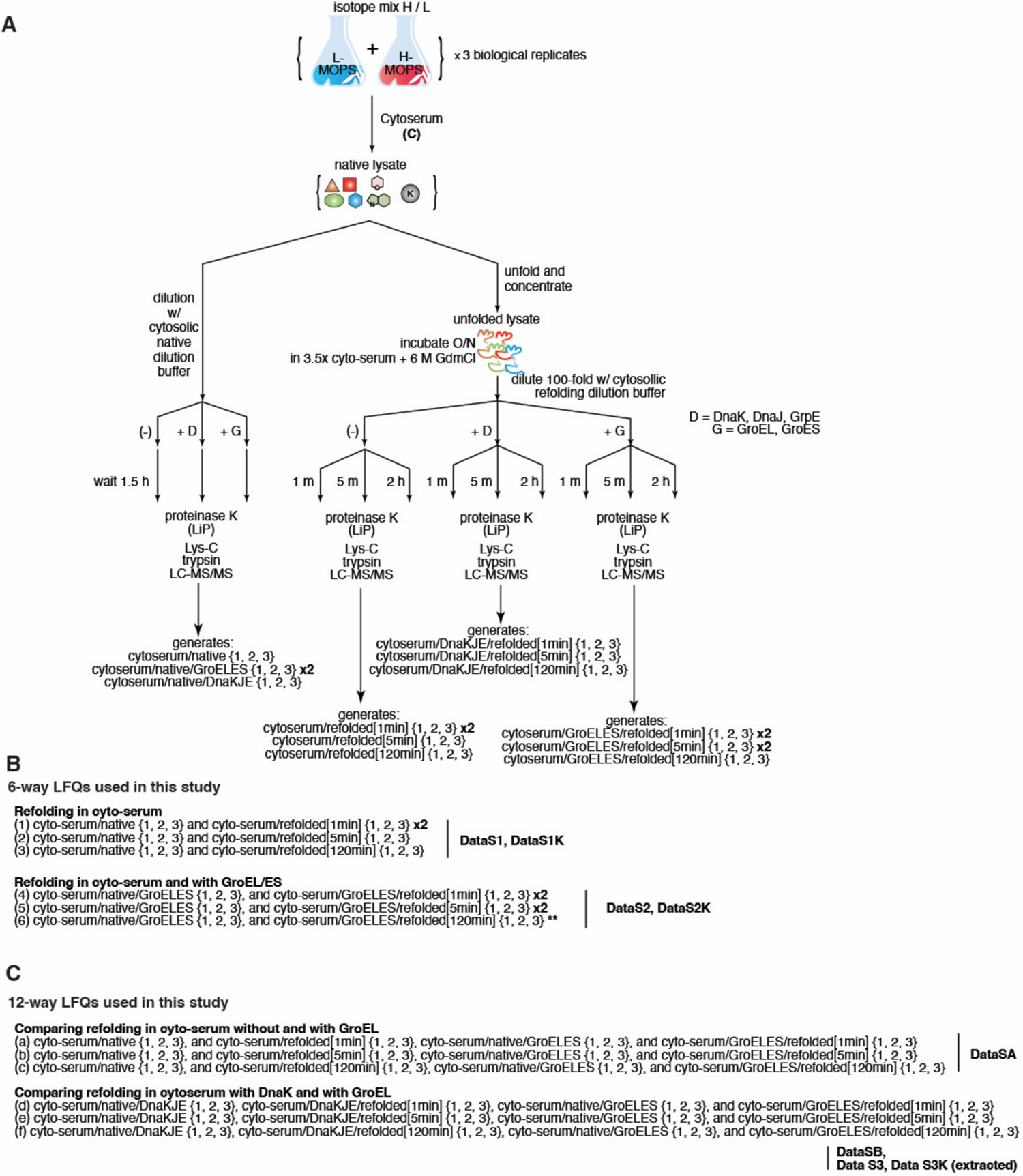
Summary of all samples prepared for LC-MS/MS, and their combinations to perform label-free quantification (LFQ) analyses. (A) Experimental workflow to prepare the 51 samples for LC-MS/MS used in the final experiments published in this study. See Materials and Methods for more details. In brief: three *E. coli* cultures are grown in light MOPS media and three *E. coli* cultures are grown in heavy MOPS media. Pairs are mixed together, and cells are gathered by centrifugation. Pellets are resuspended in cyto-serum lysis buffer. The native samples are probed by limited proteolysis (LiP) with proteinase K (PK) after equilibration. The refolded samples are probed similarly, but at 3 different timepoints following initiation of refolding by dilution (1 min, 5 min, and 120 min). The cyto-serum-lysed samples are either diluted in cyto-serum native dilution buffer to generate cyto-serum/native samples, or diluted in cyto-serum native dilution buffers supplemented with GroEL/ES or DnaK/J/E. Following equilibration, they are probed with proteinase K. Alternatively, cyto-serum lysates are unfolded into 6 M GdmCl, and refolded by 100-fold dilution into cyto-serum refolding buffer, either supplemented with GroEL/ES, DnaK/J/E, or neither, and given either 1 min, 5 min, or 120 min to refold prior to interrogation with PK. In all cases, immediately following 1 min of LiP, samples are quenched by boiling, fully trypsinized with LysC and trypsin, and prepared for LC-MS/MS. (B) Summary of the six 6-way LFQs used in this study, and which set of six samples are analyzed together to generate the peptide refolded/native quantifications. Figure 3A (1); Figure 3B (4); Figure 3C (2, 5); Figure 3D,E (2 left, 5 right); Figure 3F,G (2, 5, (Nissley et al., 2021)); Figure 5A,C,E,G (1–both reps., 2, 3); Figure 5B,D,F,H (4– both reps., 5–both reps.); Figure 5I (2, 5); Figure 6B (1–6 & see below); Figure 6C (combination of 4, 5). **The 2 h timepoint for GroEL/ES refolding was generally not used because of ATP depletion. (C) Summary of the six 12-way LFQs used in this study, and which set of twelve samples are analyzed together to generate peptide refolded/native quantifications. Figure 4A-H (b); Figure 6A (e– extracting out the DnaK subexperiment); Figure 6B (d, e, f, & see above); Figure 6D (combination of d, e); Figure 6E-H (e); Figure 7D (b).

**Figure 1– figure supplement 2.**
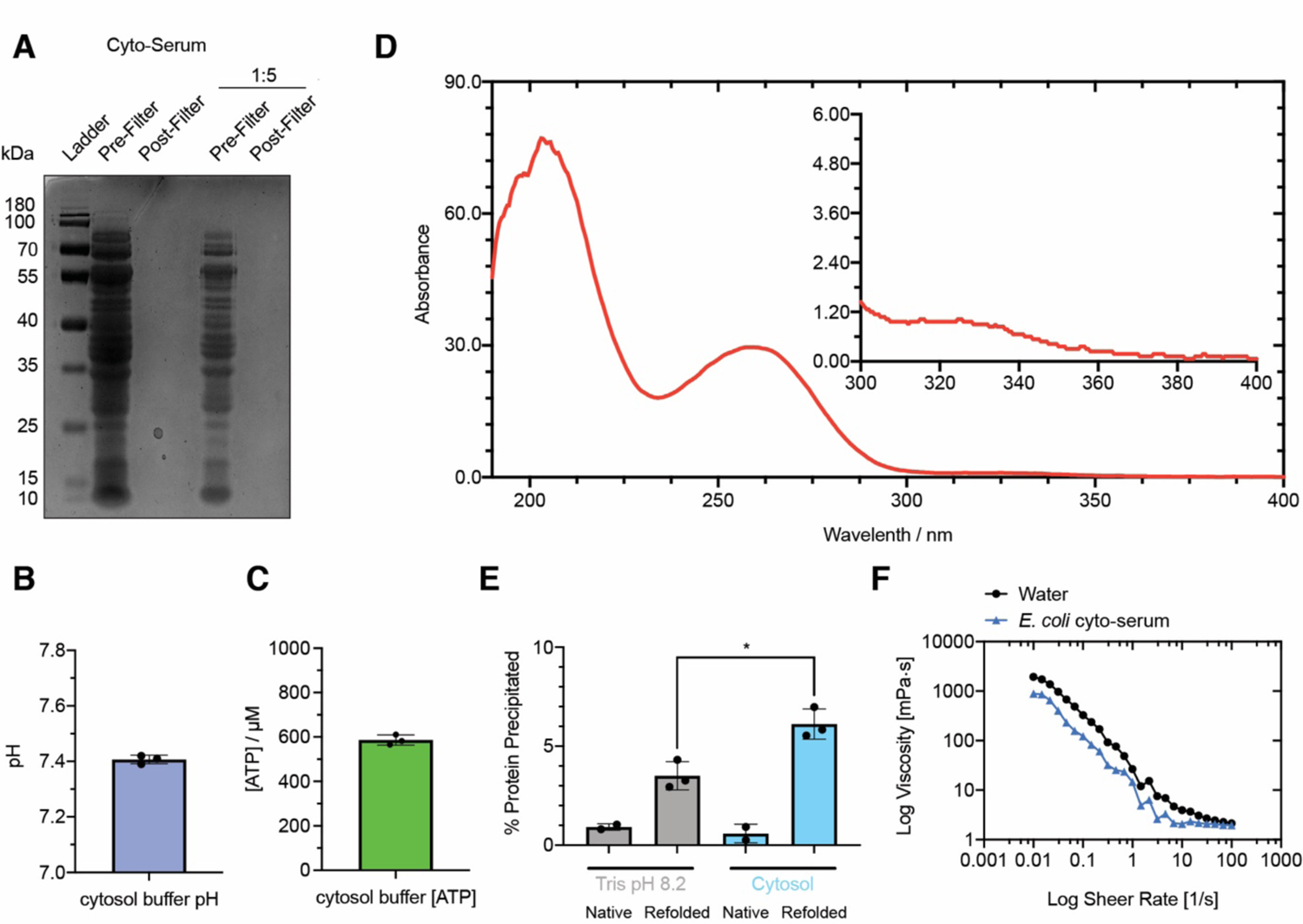
Characterization of cyto-serum. (A) Coomassie staining of SDS-PAGE of cyto-serum pre and post filtration with Viva-Spin 15R 2K MWCO Filter (Sartorius) as a 1x and a 1:5 dilution. Filtration effectively removes all macromolecules larger than 2 kDa; see experimental procedures. (B) Bar chart showing the pH readings of 3 independent preparations of cyto-serum. (C) Bar chart showing the quantification of free ATP for 3 independent preparations of cyto-serum pre-ultracentrifugation and filtration. (D) UV-VIS spectra obtained of 1x cyto-serum. Cyto-serum is an off yellow liquid with a strong absorbance at 258 nm. Absorbances observed between 200 nm – 360 nm indicate the presence of ions, metabolites, and cofactors present in cyto-serum. (E) Bar charts showing the quantification of protein aggregation of native and refolded samples in Tris pH 8.2 and cyto-serum using BCA Assay. Refolding in cyto-serum resulted in a small but significant increase in detected protein precipitation upon refolding (P < 0.05 by Welch’s t-test) when compared to refolding in Tris pH 8.2. (F) Log-log diagram showing viscosity of cyto-serum and water as a function of sheer rate. *E. coli* cyto-serum is a non-viscous fluid with rheometric properties similar to water.

**Figure 1– figure supplement 3.**
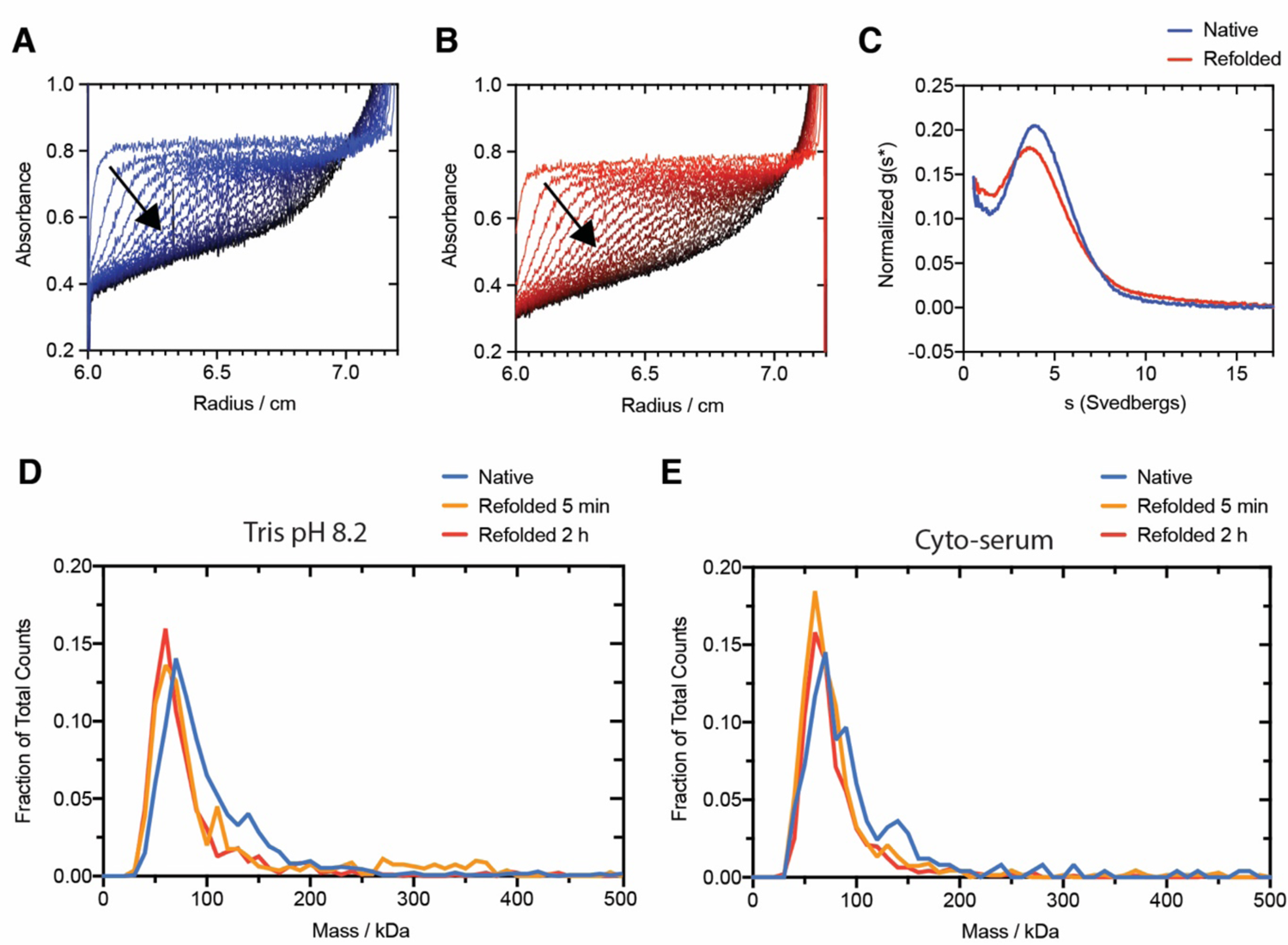
Low aggregation during global refolding reactions. (A-B) Absorbance at 280 nm as a function of radius along the rotor during sedimentation velocity analytical ultracentrifugation of native (A) and refolded (B) *E. coli* lysates in Tris pH 8.2. Data from first 100 scans are shown, with each subsequent line representing every 5^th^ scan in directionality of arrow. These datasets are representative of two independent spins on two separately prepared native and refolded lysates. (C) Calculated sedimentation coefficient distributions of native and refolded E. coli lysates in Tris pH 8.2 determined using dcdt+. Sedimentation coefficients were corrected to 20 °C in water using density, viscosity, and partial specific volume values calculated in SEDNTERP These datasets are representative of two independent spins on two separately prepared native and refolded lysates. (D-E) Normalized mass distributions of native and refolded *E. coli* lysates (5 min and 2 h) in Tris pH 8.2 (D) and cyto-serum (E) as determined by Mass Photometry (MP). All three sample types show overlaying mass distributions in both refolding buffers (Tris or cyto-serum), indicative that there are minimal differences in soluble aggregation between native and refolded samples.

**Figure 3– figure supplement 1.**
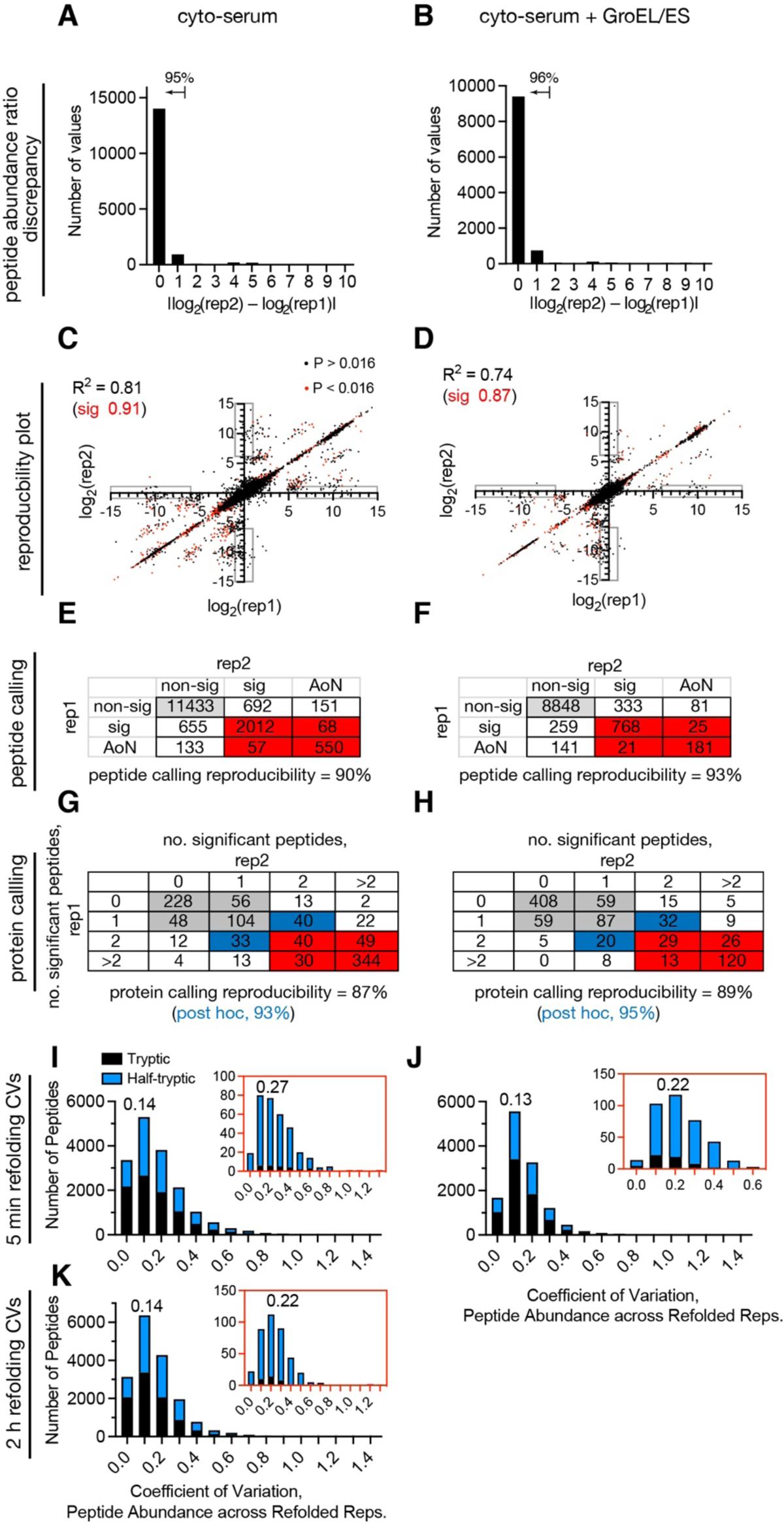
Reproducibility analysis. (A-B) Histograms showing the peptide quantification discrepancies between two replicates of the experiment in which proteins were refolded for 1 min in (A) cyto-serum, or (B) cyto-serum with GroEL/ES. These correspond to the two replicates of LFQ 1 and 4 from Figure 1–figure supplement 1B. Note that each of these replicates of the experiment involved three separate biological replicates of native and refolded. Peptides that were identified in both experiments were collected and the refolded/native ratio in each replicate was compared to each other. Histograms show the absolute value of the difference of the log2 quantifications. (A) 15751 peptides were identified in common, of which 89% were within 1.4-fold and 95% were within 3.8-fold. (B) 10564 peptides were identified in common, of which 89% were within 1.4-fold and 96% were within 3.8-fold. (C-D) Scatter plots showing the relationship between the peptide log2(refolded/native) quantification in one replicate versus its value in the other replicate for two replicates of the experiment in which proteins were refolded for 1 min in (C) cyto-serum, or (D) cyto-serum with GroEL/ES. Points in red were considered significant (P < 0.016 by Welch’s t-test) in both experiments. The coefficients of determination (*R*^2^) are given first for all points in common (black), and then for the subset of points that were considered significant in both replicates of the experiment (red). In all cases, *R*^2^ is greater when only significant peptides are considered (which are the only ones used to call a protein non-refoldable). The gray boxes demarcate regions in which upon separate performances of the experiment, an all-or-nothing peptide is categorized as nonsignificant in the other. Importantly, these boxes have very few red points. (E-F) Calling reproducibility of peptides (classified as either non-significant, significant, or all-or-nothing (AoN)) between two replicates of the experiment in which proteins were refolded for 1 min in (E) cyto-serum, or (F) cyto-serum with GroEL/ES. (G-H) Calling reproducibility of proteins between two replicates of the experiment in which proteins were refolded for 1 min in (G) cyto-serum, or (H) cyto-serum with GroEL/ES. Rows correspond to the number of peptides that were significantly different between native and refolded samples in the first replicate of the experiment, and columns correspond to the number of peptides that were significantly different in the duplication. Numbers in the table correspond to the number of proteins with that many significant peptides in each replicate. Gray cells correspond to proteins that would be called refoldable in both iterations. Red cells correspond to proteins that would be called nonrefoldable in both iterations. Cells in white would have been called differently, resulting in reproducibility from 87–89%. In all comparisons, we exclude proteins that only differ by one significant peptide at the cut-off, shown as blue cells. With these proteins removed *post hoc*, reproducibility increases to 93–95%. (I-J) Histograms of the coefficients of variation (CV) for the peptide abundances in refolded samples, from 3 independent refolding reactions, after 5 min of refolding for experiments in which cells were lysed and refolded in either (I) cyto-serum, or (J) cyto-serum with GroEL/ES. Insets in red correspond to the CV histograms for the peptides detected only in the refolded samples (which are almost all half-tryptic). Numbers represent medians of distributions. (K) Same as panels I, except for refolding after 2 h.

**Figure 3– figure supplement 2.**
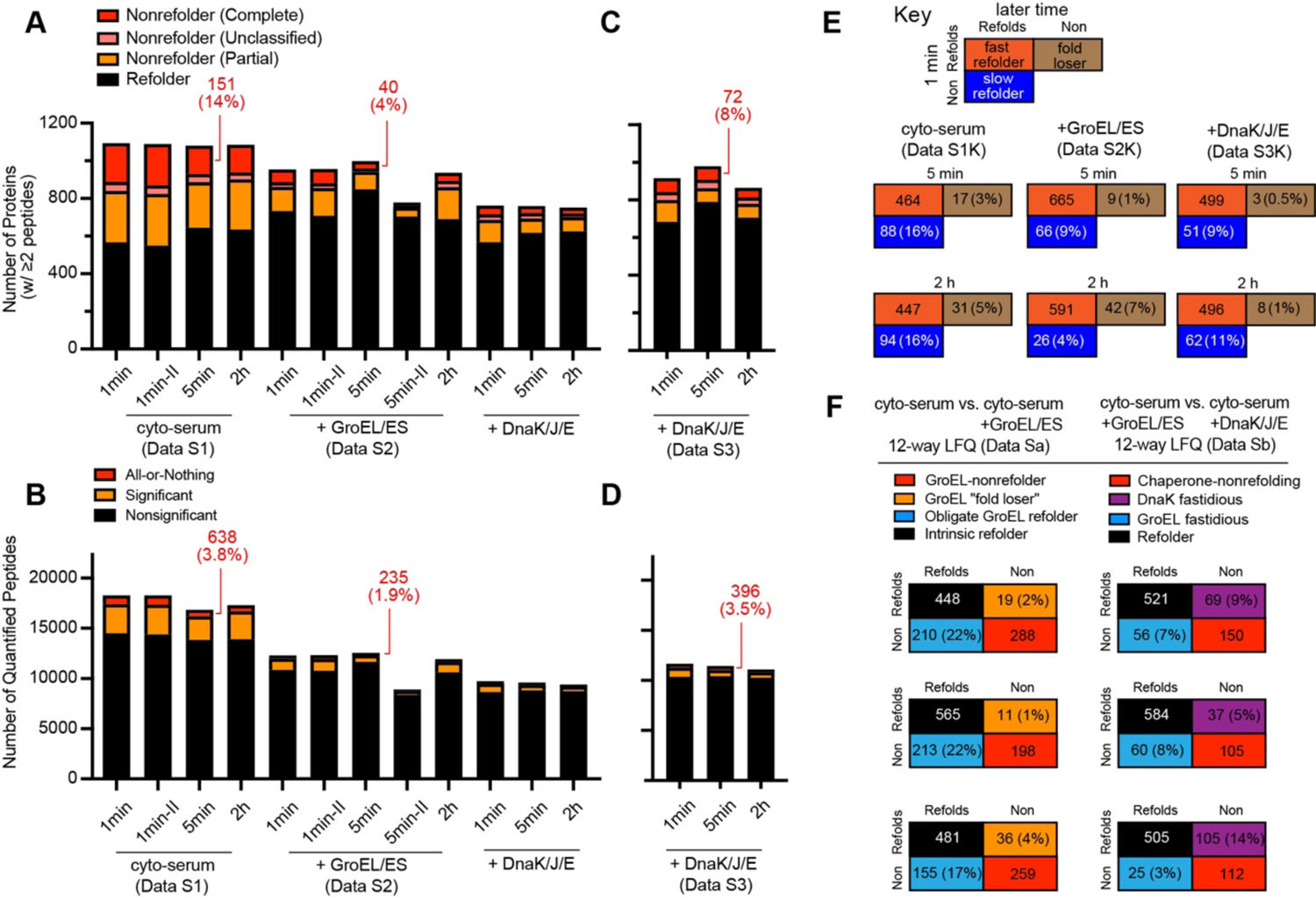
Summary statistics of all 6-way LFQs, kinetic comparisons, and truth tables for condition comparisons based on 12-way LFQs. (A) The number of proteins assessed in each 6-way LFQ, categorized as either refolding (0 or 1 peptide quantified with significantly different abundance between native and refolded), partial nonrefolding (2 or more peptides quantified with significantly different abundance between native and refolded but fewer than 2 all-or-nothing peptides), and complete nonrefolding (2 or more all-or-nothing peptides). Some nonrefolding proteins do not classify between the subcategories (if they have 1 significant and 1 all-or-nothing peptide). Proteins with only 1 peptide quantified are not included. Bars correspond in order to 6-way LFQs labeled #1–6 in Figure 1–figure supplement 1B. 6-way LFQs for DnaK/J/E are not used for analysis (see main text and methods). In red, are number (and percentage) of proteins that are judged complete nonrefolding. (B) The number of peptides confidently quantified in each 6-way LFQ, categorized as either nonsignificant, significant, or all-or-nothing. Bars correspond in order to the 6-way LFQs labeled #1–6 in Figure 1–figure supplement 1B. In red, are number (and percentage) of peptides that are all-or-nothing. (C) Data correspond to #d–f in Figure 1–figure supplement 1C, with the identifications and quantifications for the DnaK channels extracted out, done to increase coverage in the DnaK experiments (see main text). Categorizations same as panel A. (D) Data correspond to #d–f in Figure 1–figure supplement 1C, with the identifications and quantifications for the DnaK channels extracted out, done to increase coverage in the DnaK experiments (see main text). Categorizations same as panel B. (E) Summary of all kinetics experiments. To assess kinetics, we perform a comparison of two 6-way LFQs that correspond to distinct refolding timepoints but for otherwise identical conditions. To be included, a protein must have two or more confidently quantified peptides at both timepoints, be assessed as refoldable in one of the two time points, and cannot differ by only one significant peptide between the two timepoints. Each protein is designated as either a fast refolder, slow refolder, or fold loser; the number of such proteins is given for each kinetic comparison, according to the key. For the top row, from left to right, the data used for each comparison correspond to: #1 & 2; #4 & 5(Figure 1–figure supplement 1B) and #d & e (Figure 1–figure supplement 1C). For the bottom row, from left to right, the data used for each comparison correspond to #1 & 3; #4 & 6 (Figure 1– figure supplement 1B) and #d & f (Figure 1–figure supplement 1C) (F) Summary of all condition comparison experiments. To assess the effect of changing refolding condition, we perform 12-way LFQs that merge the native and refolded (at a given timepoint) replicates for the two conditions being compared. To be included, the protein must have two or more confidently quantified peptides in both conditions, and cannot differ by only one significant peptide between the two conditions. Two types of comparisons were performed (columns): cyto-serum with and without GroEL/ES, GroEL/ES vs. DnaK/J/E in cyto-serum. The color code for the designations associated with each comparison are given, and the truth tables give the number of proteins in each designation. Each comparison was conducted at three timepoints. Data correspond to #a–f in Figure 1–figure supplement 1C.

**Figure 3– figure supplement 3.**
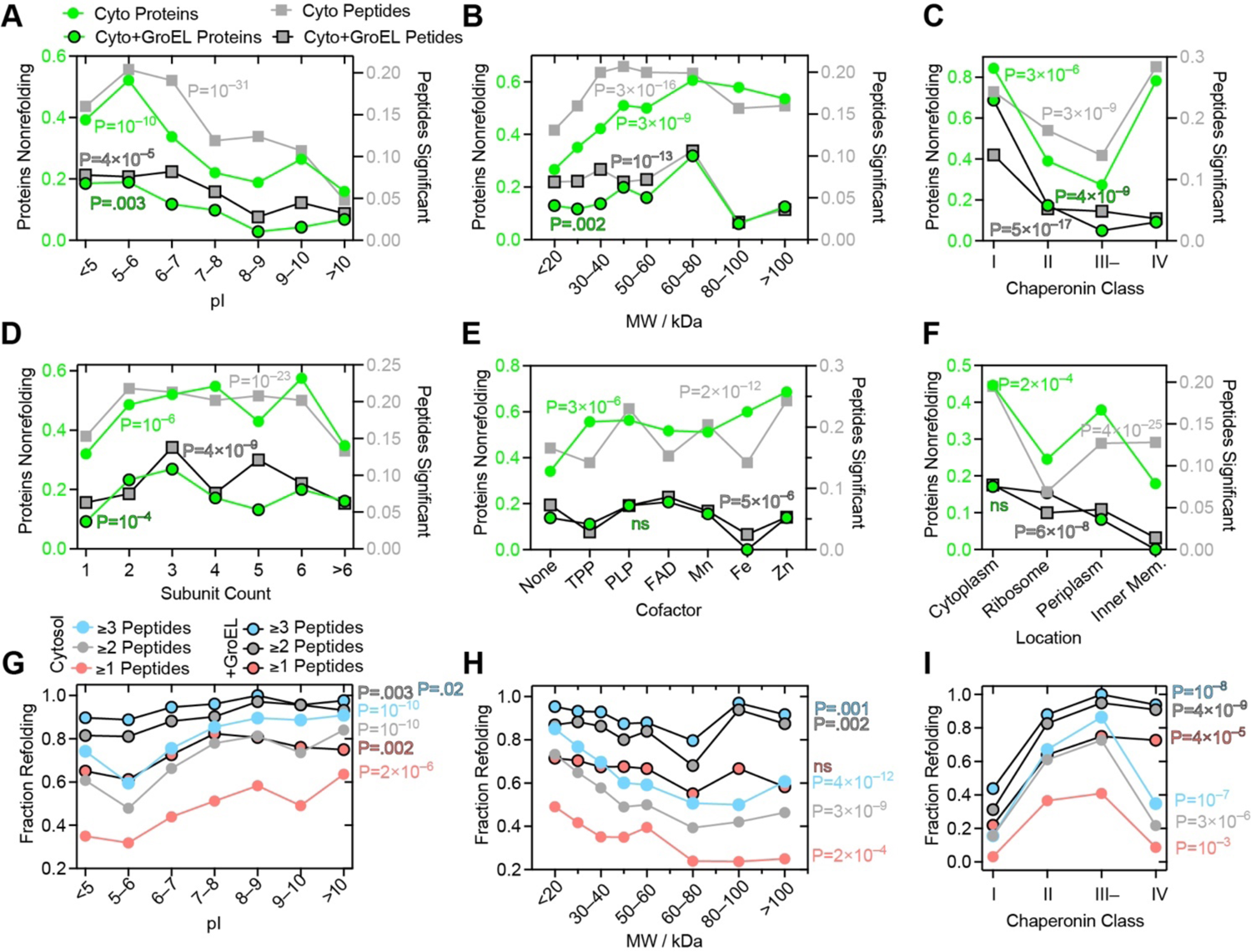
Peptide-level analyses and comparison to the protein-level. (A) Fraction of proteins that do **NOT** refold in either cyto-serum (green circles), or cyto-serum with GroEL/ES (green circles, black border), separated based on individual proteins’ isoelectric point (pI) (left y-axis). Additionally shown is fraction of peptides that have significantly different abundance in refolded samples, lumped together for all proteins within the given pI tranche (in either the cyto-serum experiment (gray boxes) or the cyto-serum with GroEL/ES experiment (gray boxes, black border)) (right y-axis). P-values according to the chi-square test are given on the protein frequencies in green (black border for GroEL/ES) and on the peptide frequencies in gray (black border for GroEL/ES). Proteins with low pI tend to be more nonrefoldable *and* tend to generate significant peptides at a much higher frequency; the trend prevails across the series with protein nonrefoldability fraction tracking closely with the peptide significance rate. This implies that the trend is *robust*, and not a coverage artefact. Data correspond to #2 and #5 in Figure 1–figure supplement 1B. (B) Same as panel A, except proteins and peptides are separated on the basis of the protein’s molecular weight (MW). Trends associated with MW are robust. (C) Same as panel A, except proteins and peptides are separated on the basis of the protein’s chaperonin class (Kerner et al., 2005). Trends associated with chaperonin class are robust. (D) Same as panel A, except proteins and peptides are separated on the basis of the protein’s subunit count. In the cytosol, monomeric and large complexes refold the most efficiently, a robust trend. With GroEL/ES, overall trends with respect to subunit count are less substantial, though it appears to have an outsized importance on tetrameric and hexameric proteins. (E) Same as panel A, except proteins and peptides are separated on the basis of the which cofactors the protein harbours. In the cytosol, apo-proteins refold the best and Fe & Zn metalloproteins refold the worst, although due to low counts the trends are not robust (the protein and peptide level results do not track together). With GroEL/ES, overall trends with respect to subunit are less substantial, though it appears to have an outsized importance holo-proteins over apo-proteins. (F) Same as panel A, except proteins are peptides are separated on the basis of the protein’s cellular localization. Trends associated with location are robust. GroEL/ES is effective on proteins in all locations, except ribosomal proteins. (G-I) Sensitivity analyses showing the fraction of proteins refolding in either cyto-serum (solid circles) or in cyto-serum with GroEL/ES (solid circles, black borders), as a function of the number of significant peptides required to call a protein nonrefoldable (≥1, red; ≥2, gray (the standard cutoff); ≥3, blue). P-values according to the chi-square test are given in matching colors (for the various cutoff schemes) and with black borders for GroEL/ES. Whilst all trends are maintained irrespective of cutoff, statistical significances generally fall with the ≥1 cutoff (red), likely because it assigns too much weight to a single significant peptide.

**Figure 6– figure supplement 1.**
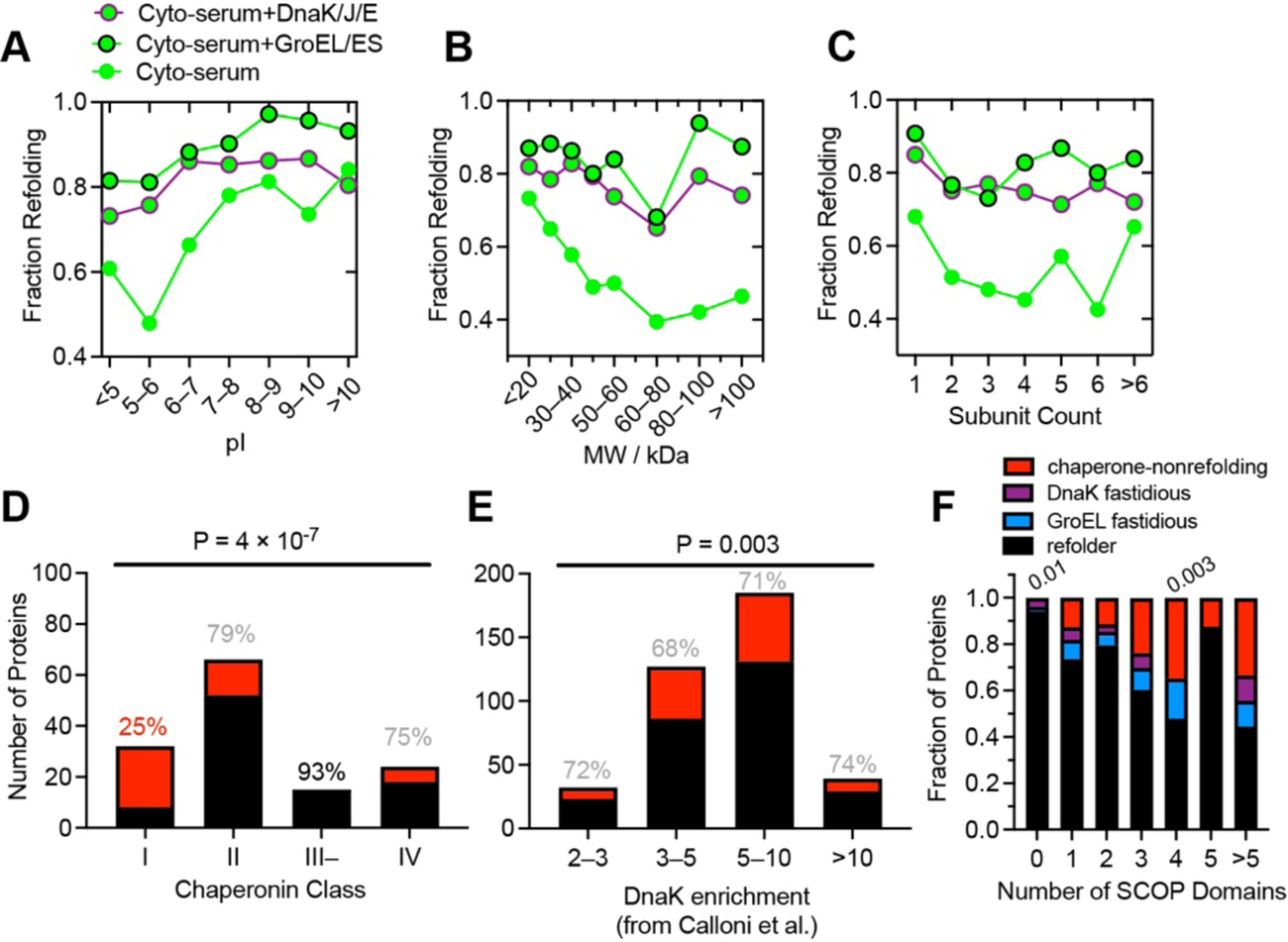
Further properties of DnaK refolders. (A) Fraction of proteins that refold in either cyto-serum (green), cyto-serum with GroEL/ES (green, black border), or cyto-serum with DnaK/J/E (green, purple borer) separated on the basis of proteins’ pI. Data come Nissley et al. (2021), #2 in Figure 1–figure supplement 1B and #e in Figure 1–figure supplement 1C. (B) As A, except proteins are separated on the basis of proteins’ molecular weight (MW). (C) As A, except proteins are separated on the basis of the number of subunits in the complex to which they are part. (D) Bar charts indicating the number of refolding (black) and nonrefolding (red) proteins associated with one of four chaperonin classes (as defined by Kerner et al., 2005; Fujiwara et al., 2010), during refolding experiments in cyto-serum with DnaK/J/E. Percents indicate percentage refolding within that category. P-value is from the chi-square test. Data come from a 12-way LFQ (#e in Figure 1–figure supplement 1C) with the identifications and quantifications from the DnaK sub-experiment extracted out. (E) As D, except proteins are separated by their enrichment level in DnaK pull-down assays (as defined by Calloni et al., 2012). (F) Frequency of proteins that refolded in both conditions (black), only with GroEL/ES (light blue), only with DnaK/J/E (purple), or did not refold in either (chaperone-nonrefolder; red), separated on the basis of the number of domains in the protein, as defined by the SCOP database. Data used for F is the same 12-way LFQ as panel D. Numbers indicate P-values according to the chi-square test.

## REFERENCES

1. Ambrose, A. J., Fenton, W., Mason, D. J., Chapman, E., Horwich, A. L. (2015). Unfolded DapA forms aggregates when diluted into free solution, confounding comparison with folding by the GroEL/GroES chaperonin system. FEBS Lett. 589, 497–499.

2. Anfinsen, C. B., Haber, E., Sela, M., and White Jr., F. H. (1961). The kinetics of formation of native ribonuclease during oxidation of the reduced polypeptide chain. Proc. Natl. Acad. Sci. USA 47, 1309–1314.

3. Anfinsen, C. B. (1973). Principles that govern the folding of protein chains. Science 181, 223– 230.

4. Balchin, D., Hayer-Hartl, M., and Hartl, F. U. (2016). In vivo aspects of protein folding and quality control. Science 353, aac4354.

5. Balchin, D., Hayer-Hartl, M., and Hartl F. U. (2020). Recent advances in understanding catalysis of protein folding by molecular chaperones. FEBS Lett. 594, 2770–2781.

6. Bartlett AI and Radford SE. (2009). An expanding arsenal of experimental methods yields an explosion of insights into protein folding mechanisms. Nat Struct Mol Biol 16, 582–588.

7. Bennett, B. D., Kimball, E. H., Gao, M., Osterhout, R., van Dien, S. J., and Rabinowitz, J. D. (2009). Absolute metabolite concentrations and implied enzyme active site occupancy in Escherichia coli. Nature Chem. Biol. 5, 593–599.

8. Bertolini, M.,… and Kramer, G. (2021). Interactions between nascent proteins translated by adjacent ribosomes drive homomer assembly. Science 371, 57–64.

9. Bowman, J. C., Petrov, A. S., Frenkel-Pinter, M., Penev, P. I., and Williams, L. D. (2020). Root of the Tree: The Significance, Evolution, and Origins of the Ribosome. Chem. Rev. 120, 4848– 4878.

10. Brinker, A… (2001). Dual Function of Protein Confinement in Chaperonin-Assisted Protein Folding. Cell 107, 223–233.

11. Chaudhuri, T. K., Farr, G. W., Fenton, W. A., Rospert, S., and Horwich A. L. (2001). GroEL/GroES-Mediated Foldingof a Protein Too Largeto Be Encapsulated. Cell 107, 235–246.

12. Chaudhuri, T. K., Verma, V. K., and Maheshwari, A. (2009). GroEL assisted folding of large polypeptide substrates in *Escherichia coli*: Present scenario and assignments for the future. Prog. Biophys. Mol. Biol. 99, 42–50.

13. Calloni, G., Chen, T., Schermann, S. M., Chang, H., Genevaux, P., Agostini, F., Tartaglia, G. G., Hayer-Hartl, M., and Hartl, F. U. (2012). DnaK Functions as a Central Hub in the E. coli Chaperone Network. Cell Reports 1, 251–264.

14. Cheng, H., et al. (2014). ECOD: An Evolutionary Classification of Protein Domains. PLOS Comput. Biol. 10, e1003926.

15. Cole, D., Young, G., Weigel, A., Sebesta, A., and Kukura, P. (2017). Label-Free Single-Molecule Imaging with Numerical-Aperture-Shaped Interferometric Scattering Microscopy. ACS Photonics 4, 211–216.

16. De Souza, N., and Picotti, P. (2020). Mass spectrometry analysis of the structural proteome. Curr. Opin. Struct. Biol. 60, 57–65.

17. Farr, G. W., Fenton, W. A., Chaudhuri, T. K., Clare, D. K., Saibil, H. R., and Horwich, A. L. (2003). Folding with and without encapsulation by *cis*- and *trans*-only GroEL–GroES complexes. EMBO J. 22, 3220–3230.

18. Feng, Y., Franceschi, G., Kahraman, A., Soste, M., Melnik, A., Boersema, P. J., de Laureto, P. P., Nikolaev, Y., Oliveria, A. P., and Picotti, P. (2014). Global analysis of protein structural changes in complex proteomes. Nature Biotech. 32, 1036–1044.

19. Fedorov, A. N. and Baldwin, T. O. (1997). Cotranslational protein folding. J. Biol. Chem. 272, 32715–32718.

20. Fried, S. D., Fujishima, K., Makarov, M., Cherepashuk, I., and Hlouchova, K. (2022). Peptides Before and During the Nucleotide World: An Origins Story Emphasizing Cooperation between Proteins and Nucleic Acids. J. Roy. Soc. Interface 19. DOI: 10.1098/rsif.2021.0641.

21. Frydman, J., Erdjument-Bromage, H., Tempst, P., and Hartl, F. U. (1999). Co-translational domain folding as the structural basis for the rapid de novo folding of firefly luciferase. Nature Struct. Biol. 6, 697–705.

22. Fujiwara, K., Ishihama, Y., Nakahigashi, K., Soga, T., and Taguchi, H. (2010). A systematic survey of *in vivo* obligate chaperonin-dependent substrates. EMBO J. 29, 1552–1564.

23. Gao, X., et al. (2015). Human Hsp70 Disaggregase Reverses Parkinson’s-Linked α-Synuclein Amyloid Fibrils. Mol. Cell. 59, 781–793.

24. Georgescauld, F., Popova, K., Gupta, A. J., Bracher, A., Engen, J. R., Hayer-Hartl, M., and Hartl, F. U. (2014). GroEL/ES Chaperonin Modulates the Mechanism and Accelerates the Rate of TIM-Barrel Domain Folding. Cell 157, 922–934.

25. Goloubinoff, P., Christeller, J. T., Gatenby, A. A., and Lorimer, G. H. (1989). Reconstitution of active dimeric ribulose bisphosphate carboxylase from an unfolded state depends on two chaperonin proteins and Mg-ATP. Nature 342, 884–889.

26. Gottesman, S., Wickner, S., and Maurizi, M. R. (1997). Protein quality control: triage by chaperones and proteases. Genes Dev. 11, 815–823.

27. Gough, J., Karplus, K., Hughey, R., and Chothia, C. (2001) Assignment of homology to genome sequences using a library of hidden Markov models that represent all proteins of known structure. J. Mol. Biol. 313, 903– 919.

28. Halder, R., Nissley, D. A., Sitarik, I., and O’Brien E. P. (2021). Subpopulations of soluble, misfolded proteins commonly bypass chaperones: How it happens at the molecular level. bioRxiv 10.1101/2021.08.18.456736.

29. Han, J.-H., Batey, S., Nickson, A. A., Teichmann, S. A., and Clarke, J. (2007). The folding and evolution of multidomain proteins. Nature Rev. Mol. Cell. Biol. 8, 319–330.

30. Harris, C. L. (1987). An aminoacyl-tRNA synthetase complex in *Escherichia coli*. J. Bacteriol. 169, 2718– 2723.

31. Houry, W. A., Frishman, D., Eckerskorn, C., Lottspeich, F., and Hartl, F. U. (1999). Identification of in vivo substrates of the chaperonin GroEL. Nature 402, 147–154.

32. Imamoglu, R., Balchin, D., Hayer-Hartl, M., and Hartl, F. U. (2020). Bacterial Hsp70 resolves misfolded states and accelerates productive folding of a multi-domain protein. Nature Comm. 11, 365.

33. Jarzab, A., et al. (2020). Meltome atlas—thermal proteome stability across the tree of life. Nature Meth. 17, 495–503.

34. Jumper, J., et al. (2021). Highly accurate protein structure prediction with AlphaFold. Nature 596, 583–589.

35. Kerner, M. J., et al. (2005). Proteome-wide Analysis of Chaperonin-Dependent Protein Folding in *Escherichia coli*. Cell 122, 209–220.

36. Keseler, I. M., et al. (2017). The EcoCyc database: reflecting new knowledge about *Escherichia coli* K-12. Nucleic Acids Res. 45, D543–D550.

37. Kong, A. T., Leprevost, F. V., Avtonomov, D. M., Mellacheruvu, D., and Nesvizhskii, A. I. (2017). MSFragger: ultrafast and comprehensive peptide identification in mass spectrometry-based proteomics. Nature Meth. 14, 513–520.

38. Kozlowski, L. P. (2017). Proteome-pI: proteome isoelectric point database. Nucleic Acids Res. 45, D1112–D1116.

39. Langer, T., Lu, C., Echols, H., Flanagan, J., Hayer, M. K., and Hartl, F. U. (1992). Successive action of DnaK, DnaJ and GroEL along the pathway of chaperone-mediated protein folding. Nature 356, 683–689.

40. Laue, T. M., Shah, B. D., Ridgeway, T. M., and Pelletier, S. L. (1992). Computer-aided interpretation of analytical sedimentation data for proteins. In: Harding, S.; Rowe, A.; Hoarton, J., editors. Analytical Ultracentrifugation in Biochemistry and Polymer Science. Royal Society of Chemistry; Cambridge, UK: 1992. p. 90-125.

41. Li, G.-W., Burkhardt, D., Gross, C., and Weissman, J. S. (2014). Quantifying absolute protein synthesis rates reveals principles underlying allocation of cellular resources. Cell 157, 624– 635.

42. Lin, Z., Madan, D., and Rye, H. S. (2008). GroEL stimulates protein folding through forced unfolding. Nature Struct. Mol. Biol. 15, 303–311.

43. Lindquist, S. (2009). Protein folding sculpting evolutionary change. Cold Spring Harb. Symp. Quant. Biol. 74, 103–108.

44. Liu, K., Maciuba, K., and Kaiser C. M. (2019). The Ribosome Cooperates with a Chaperone to Guide Multi-domain Protein Folding. Mol. Cell. 74, 310–319.

45. Macošek, J., Mas, G., and Hiller, S. (2021). Redefining Molecular Chaperones as Chaotropes. Frontiers Mol. Biosci. 8, 683132.

46. Mateus, A., Bobonis, J., Kurzawa, N., Stein, F., Helm, D., Hevler, J., Typas, A., and Savitski, M. M. (2018). Thermal proteome profiling in bacteria: probing protein state *in vivo*. Mol. Sys. Biol. 14, e8242.

47. Mayer, M. P. & Gierasch, L. M. Recent advances in the structural and mechanistic aspects of Hsp70 molecular chaperones. J. Biol. Chem. 294, 2085–2097 (2019).

48. Mi, H., Muruganujan, A., Huang, J. X., Ebert, D., Mills, C., Guo, X., and Thomas, P. D. (2019). Protocol Update for large-scale genome and gene function analysis with the PANTHER classification system (v.14.0). Nature Protoc. 14, 703–721.

49. Nahnsen, S., Bielow, C., Reinert, K., and Kohlbacher, O. (2013). Tools for label-free peptide quantification. Mol. Cell. Proteomics 12, 549–556.

50. Neidhardt, F. C., Bloch, P. L., and Smith, D. F. (1974). Culture medium for enterobacteria. J. Bacteriol. 119, 736–747.

51. Nissley, D., Jiang, Y., Trovato, F., Sitarik, I., Narayan, K., To, P., Xia, Y., Fried, S. D., and O’Brien, E. P. (2021). Universal protein misfolding intermediates can bypass the proteostasis network and remain soluble and non-functional. bioRxiv 10.1101/2021.08.18.456613.

52. Niwa, T., Ying, B.-W., Saito, K., Jin, W., Takada, S., Ueda, T., and Taguchi, H. (2009). Bimodal protein solubility distribution revealed by an aggregation analysis of the entire ensemble of Escherichia coli proteins. Proc. Natl. Acad. Sci. USA 106, 4201–4206.

53. Niwa, T., Kanamori, T., Ueda, T., and Taguchi, H. (2012). Global analysis of chaperone effects using a reconstituted cell-free translation system. Proc. Natl. Acad. Sci. USA 109, 8937–8942.

54. Palomba, A., Abbondio, M., Friorito, G., Uzzau, S., Pagnozzi, D., and Tanca, A. (2021). Comparative Evaluation of MaxQuant and Proteome Discoverer MS1-Based Protein Quantification Tools. J. Proteome. Res. 20, 3497–3507.

55. Pandurangan, A. P., Stahlacke, J., Oates, M. E., Smithers, B., and Gough, J. (2019). The superfamily 2.0 database: A significant proteome update and a new webserver. Nucleic Acids Res. 47, D490– D494.

56. Park, C., and Marqusee, S. (2005). Pulse proteolysis: A simple method for quantitative determination of protein stability and ligand binding. Nature Meth. 2, 207–212.

57. Park, C., Zhou, S., Gilmore, J., and Marqusee, S. (2007). Energetics-based Protein Profiling on a Proteomic Scale: Identification of Proteins Resistant to Proteolysis. J. Mol. Biol. 368, 1426–1437.

58. Paul, S., Singh, C., Mishra, S., and Chaudhuri, T. K. (2007). The 69 kDa *Escherichia coli* maltodextrin glucosidase does not get encapsulated underneath GroES and folds through *trans* mechanism during GroEL/ GroES-assisted folding. FASEB J. 21, 2874–2885.

59. Petrov, A.S. et al. (2015). History of the ribosome and the origin of translation. Proc. Natl. Acad. Sci. USA 112, 15396–15401.

60. Phillips, R., Kondev, J., and Theriot, J. (2008) Physical Biology of the Cell. Garland Science, New York, NY.

61. Philo, J. S. (2006). Anal Biochem. 354, 238–246.

62. Powers, E. T., Powers, D. L., and Gierasch, L. M. (2012). FoldEco: A Model for Proteostasis in *E. coli*. Cell Rep. 1, 265–276.

63. Rebeaud, M. E., Mallik, S., Goloubinoff, P., and Tawfik, D. S. (2021). On the evolution of chaperones and co-chaperones and the expansion of proteomes across the Tree of Life. Proc. Natl. Acad. Sci. USA 118, e2020885118.

64. Rosenzweig, R., Nillegoda, N. B., Mayer, M. P. & Bukau, B. The Hsp70 chaperone network. Nat. Rev. Mol. Cell Biol. 20, 665–680 (2019).

65. Santra, M., Farrell, D. W., and Dill, K. A. (2017). Bacterial proteostasis balances energy and chaperone utilization efficiently. Proc. Natl. Acad. Sci. USA *X*, E2654–E2661.

66. Schaefer, R. D., Liao, Y., Cheng, H., and Grishin, N. V. (2016). ECOD: new developments in the evolutionary classification of domains. Nucleic Acids Res. 45, D296–D302.

67. Shiber, A., et al. (2018). Cotranslational assembly of protein complexes in eukaryotes revealed by ribosome profiling. Nature 561, 268–272.

68. Shieh, Y.-W., Minguez, P., Bork, P., Auburger, J. J., Guilbride, D. L., Kramer, G., and Bukau, B. (2015). Operon structure and cotranslational subunit association direct protein assembly in bacteria. Science 350, 678–680.

69. Singh, A. K., Balchin, D., Immamoglu, R., Hayer-Hartl, M., and Hartl, F. U. (2020). Efficient Catalysis of Protein Folding by GroEL/ES of the Obligate Chaperonin Substrate MetF. J. Mol. Biol. 432, 2304–2318.

70. Tang, Y.-C., Chang, H.-C., Roeben, A., Wichnewski, D., Wichnewski, N., Kerner, M. J., Hartl, F. U., Hayer-Hartl, M. (2006). Structural features of the GroEL-GroES nano-cage required for rapid folding of encapsulated protein. Cell 125, 903–914.

71. Thirumalai, D., and Lorimer, G. H. (2001). Chaperonin-mediated protein folding. Annu. Rev. Biophys. 30, 245–269.

72. To, P., Whitehead, B., Tarbox, H. E., and Fried, S. D. (2021). Nonrefoldability is Pervasive Across the *E. coli* Proteome. J. Am. Chem. Soc. 143, 11435–11448.

73. Tyedmers, J., Mogk, A., and Bukau, B. (2010). Cellular strategies for controlling protein aggregation. Nature Rev. Mol. Cell Biol. 11, 777–788.

74. Vecchi, G., Sormanni, P., Mannini, B., Vandelli, A., Tartaglia, G. G., Dobson, C. M., Hartl, F. U., and Vendruscolo, M. (2020). Proteome-wide observation of the phenomenon of life on the edge of solubility. Proc. Natl. Acad. Sci. USA 117, 1015–1020.

75. Viitanen, P. V., Lubben, T. H., Reed, J., Goloubinoff, P., O’Keefe, D. P., and Lorimer, G. H. (1990). Chaperonin-Facilitated Refolding of Ribulosebisphosphate Carboxylase and ATP Hydrolysis by Chaperonin 60 (groEL) Are K+ Dependent. Biochemistry 29, 5665–5671.

76. Wallace, E. W. J., et al. (2015). Reversible, Specific, Active Aggregates of Endogenous Proteins Assemble upon Heat Stress. Cell 162, 1286–1298.

77. Weissman, J. S., Rye, H. S., Fenton, W. A., Beecham, J. M., and Horwich, A. L. (1996) Characterization of the active intermediate of a GroEL-GroES-mediated protein folding reaction. Cell 84, 481–490.

78. Willmund, F., del Alamo, M., Pechmann, S., Chen, T., Albanèse, V., Dammer, E. B., Peng, J., and Frydman, J. (2013). The cotranslational function of ribosome-associated Hsp70 in eukaryotic protein homeostasis. Cell 152, 196–209.

79. Yam, A. Y., Xia, Y., Lin H.-T. J., Burlingame, A., Gerstein, M., and Frydman, J. (2008). Defining the TRiC/CCT interactome links chaperonin function to stabilization of newly made proteins with complex topologies. Nature Struct. Mol. Biol. 15, 1255–1262.

80. Ying, B.-W., Taguchi, H., Kondo, M., and Ueda, T. (2005). Co-translational Involvement of the Chaperonin GroEL in the Folding of Newly Translated Polypeptides. J. Biol. Chem. 280, 12035–12040.

81. Young, G., et al. (2018). Quantitative mass imaging of single biological macromolecules. Science 360, 423–427.

